# Enhancing motor learning by increasing stability of newly formed dendritic spines in motor cortex

**DOI:** 10.1101/2021.01.27.428554

**Authors:** Eddy Albarran, Aram Raissi, Omar Jáidar, Carla J. Shatz, Jun B. Ding

## Abstract

Dendritic spine dynamics of Layer 5 Pyramidal neurons (L5PNs) are thought to be physical substrates for motor learning and memory of motor skills and altered spine dynamics are frequently correlated with poor motor performance. Here we describe an exception to this rule by studying mice lacking Paired immunoglobulin receptor B (PirB−/−). Using chronic two-photon imaging of primary motor cortex (M1) of PirB−/−;Thy1-YFP-H mice, we found a significant increase in the survival of spines on apical dendritic tufts of L5PNs, as well as increased spine formation rates and spine density. Surprisingly and contrary to expectations, adult PirB−/− mice learn a skilled reaching task more rapidly compared to wild type (WT) littermate controls. Conditional excision of PirB from forebrain pyramidal neurons in adult mice replicated these results. Furthermore, chronic imaging of L5PN dendrites throughout the learning period revealed that the stabilization of learning-induced newly formed spines is significantly elevated in PirB−/− mice. The degree of survival of newly formed spines in M1 yielded the strongest correlation with task performance, suggesting that this increased spine stability is advantageous and can translate into enhanced acquisition and maintenance of motor skills. Notably, inhibiting PirB function acutely in M1 of adult WT mice throughout training increases the survival of spines formed during early training and enhances motor learning. These results suggest that increasing the stability of newly formed spines is sufficient to improve long-lasting learning and motor performance and demonstrate that there are limits on motor learning that can be lifted by manipulating PirB, even in adulthood.

## INTRODUCTION

Throughout life, animals have an incredible capacity for learning new motor skills. Is this capacity optimized fully, or might there be limits on motor learning that could be removed to yield even better performance? One prevailing idea is that the acquisition of new skills involves the formation and stabilization of new dendritic spines on pyramidal cells in motor cortex (Xu et al., 2009; Yang et al., 2009; Peters et al., 2017).

Structural plasticity, the dynamic changes in spine size, shape, and numbers, is known to accompany experience-dependent changes in circuits (Yuste et al., 2001; Froemke et al., 2007 Yasumatsu et al., 2008; Holtmaat and Svoboda, 2009). For example, in the visual system, unilateral eye closure increases the rate of spine formation and leads to a lasting increase in spine density (Hofer et al., 2009); in the somatosensory system, experience drives the formation and elimination of synapses (Trachtenberg et al., 2002; Zuo et al., 2005). In the motor system, new spines on apical dendrites of L5PNs are generated when new motor skills are learned (Harms et al., 2008; Xu et al., 2009). These new experience-induced spines represent structural and functional circuit rewiring. The appearance and maintenance of these new spines involve specific changes in spine dynamics including stabilization, elimination, and turnover. In motor cortex, this increase in new spine formation on L5PNs is also accompanied by increased elimination of other spines. It is thought that learning-induced spine formation and stabilization underlie the resulting improved performance on motor tasks. This conclusion is based on several important experiments. Firstly, spine formation and maintenance, as well as spine elimination, are correlated with motor performance (Xu et al., 2009; Yang et al., 2009). Secondly, animal models of pathological conditions such as Alzheimer’s (Tsai et al., 2004; Spires et al., 2005; Shankar et al., 2007) and Parkinson’s disease (Stephens et al., 2005; Guo et al., 2015) exhibit spine destabilization and/or an acceleration of spine loss correlating with impaired learning and memory loss. Finally, directly eliminating newly formed task-specific spines abolishes learning and memory in motor cortex (Hayashi-Takagi et al., 2015; Frank et al., 2018).

All these experiments mentioned above involve manipulations or disease models that destabilize and/or eliminate spines, leading to impaired learning and memory. Conversely, there are very few studies examining the effects of increasing spine stability and/or density. In Fragile-X syndrome mice, spine density is increased but learning is impaired, suggesting that this change is detrimental (Padmashri et al., 2013). In contrast, enhanced learning on hippocampal memory tasks is associated with an increase in the number of clustered spines in mice lacking CCR5 (Frank et al., 2018). Together, these observations suggest that spine density alone cannot predict learning and memory performance and raise the possibility that spine dynamics, namely the balance between elimination and maintenance, may be a key determinant of behavioral performance.

Here we investigate this possibility by studying mice lacking PirB, a receptor expressed on pyramidal neurons throughout neocortex and hippocampus. Germline (Syken et al, 2006) or conditional deletion of PirB exclusively from neurons (Bochner et al., 2014; Djurisic et al., 2019) results in enhanced synaptic and circuit plasticity in visual cortex associated with significantly higher spine density along the dendrites of L2/3 and L5 pyramidal neurons (Djurisic et al., 2013; Vidal et al., 2016). Thus, PirB−/− mice offer an opportunity to examine the relationship between learning, increased spine density and synaptic plasticity in the motor system. To explore the consequences of PirB deletion on motor learning, mice were trained on a single-pellet reaching task and performance was assessed. The number of dendritic spines and their dynamics were measured using chronic *in vivo* two-photon imaging in M1 of PirB−/−;Thy1-YFP-H mice. To relate structural plasticity to functional plasticity, Hebbian synaptic plasticity of L5PNs was assessed in slices of M1 *in vitro*. Finally, PirB function was acutely blocked by infusing a decoy PirB receptor into M1 of adult wild type mice via minipumps to determine if the superior motor learning observed in PirB−/− mice is due to developmental compensation or to an ongoing function of PirB throughout life. Together our observations identify the stabilization of newly formed spines as a critical determinant of acquisition and maintenance of motor memory and show that there is a PirB-dependent limit that can be lifted to generate superior motor learning even in adulthood.

## RESULTS

### Increased Spine Density and Functional Synapses in M1 L5 Pyramidal Neurons of PirB−/− Mice

To investigate the impact of PirB function on excitatory synapses in motor cortex *in vivo*, two-photon microscopy was used to image spines along the apical dendrites of L5PNs through cranial windows chronically implanted over the forelimb area of M1. L5PNs were labeled by expression of yellow fluorescent protein in Thy1-YFP-H;PirB−/− mice or Thy1-YFP-H;PirB+/+ wildtype (WT) littermate controls. Two-weeks following cranial window implantation surgery, the same apical dendrites of L5PNs were imaged throughout a 6-day period at two-day intervals, allowing us to quantify formation of new spines or elimination of existing ones (Figures 1A and 1B). The rate of spine formation but not elimination was significantly greater (~25%) in PirB−/− mice compared to WT littermates (Figures 1C,D and S1A), resulting in greater spine density in PirB−/− (~31%; Figure 1E). The elevated spine density on L5PNs in PirB−/− motor cortex is consistent with previous observations in visual cortex (Djurisic et al., 2013; Bochner et al, 2014), where it was shown to arise as a consequence of deficient pruning during development (Vidal et al, 2016). Next, the survival of pre-existing and newly formed dendritic spines was monitored over the 6 day period. The survival of both pre-existing and newly formed spines was greater in PirB−/− mice compared to WT controls (Figures 1F,G). Furthermore, when spine density changes are normalized to the density measured at the first imaging session, L5PNs in PirB−/− mice have significantly lower net spine loss over days compared to WT (Figure S1B), supporting the idea that the increased spine survival and rate of formation result in the greater spine density observed in PirB−/−. When clustering of spines along dendrites is compared between WT and PirB−/− (Figures S1C,D), PirB−/− mice have a greater proportion of spines located within 0.2-2.0 μm of each other, reflecting an even greater level of spine clustering than WT.

**Figure 1.**
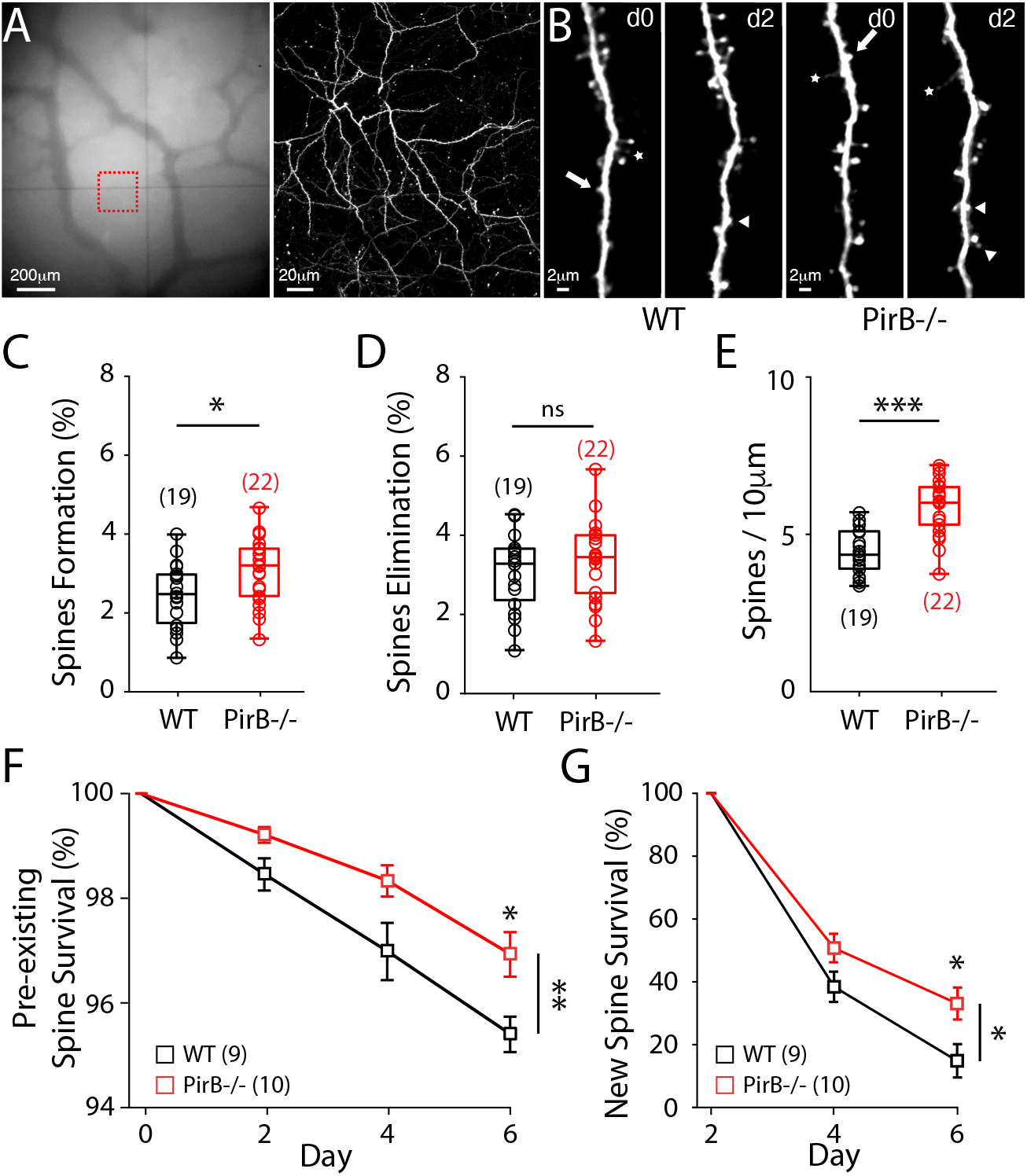
Chronic 2P Imaging of L5 Pyramidal Neuron Dendritic Spines in WT and PirB−/− Mice. (A) Repeated imaging of dendritic spines (2-day interval) in 2-month old WT and PirB−/− mice. Left: vasculature in M1 under the cranial window of a Thy1-YFP-H control mouse. Right: two photon z-projection of red square region from left, revealing apical dendrites of layer V pyramidal neurons. (B) Representative dendrites depicting spine formation and spine elimination across 2 days in WT and PirB−/− mice. Arrows depict spines to be eliminated, arrow heads depict newly formed spines. (C-D) Increased baseline rate of spine formation (WT: 2.4445 ± 0.1856%; n = 19 mice; PirB−/−: 3.0625 ± 0.1732%; n = 22 mice; p = 0.024, Mann-Whitney), but not spine elimination (WT: 3.0544 ± 0.2175%; PirB−/−: 3.3660 ± 0.2112%; p = 0.353, Mann-Whitney). (E) Density of dendritic spines was increased in PirB−/− mice (WT: 4.451 ± 0.1610; n = 19 mice; PirB−/−: 5.846 ± 0.1870; n = 22 mice; p = 1.7022 e-05, Mann-Whitney). (F) Pre-existing spine survival, quantified as the % of spines identified on the first imaging session that were present in subsequent imaging sessions. Greater survival of pre-existing spines in PirB−/− mice compared to WT (WT: n = 9; PirB−/−: n = 10; p = 0.0093, 2-way repeated measures ANOVA). Individual day statistics (day 2, WT: 98.47 ± 0.31%, PirB−/−: 99.22 ± 0.15%, p = 0.0653; day 4, WT: 97.01 ± 0.54%, PirB−/−: 98.35 ± 0.30%, p = 0.0789; day 6, WT: 95.43 ± 0.34%, PirB−/−: 96.95 ± 0.42%, p = 0.022, Mann-Whitney). (G) New spine survival, quantified as the % of spines identified on the second imaging session that were later present in subsequent imaging sessions. Greater survival of new spines in PirB−/− mice compared to WT (WT: n = 9; PirB−/−: n = 10; p = 0.031, 2-way repeated measures ANOVA). Individual day statistics (day 4, WT: 38.42 ± 4.82%, PirB−/−: 50.76 ± 4.54%, p = 0.1273; day 6, WT: 14.86 ± 5.31%, PirB−/−: 33.10 ± 5.09%, p = 0.0499, Mann-Whitney). (C-E) Circles represent individual mice. Box plots and mean ± s.e.m. are shown. (F, G) Squares depict mean survival % (± s.e.m.) across animals within each genotype. (C-G) * p < 0.05, ** p < 0.01, *** p < 0.001; ns: non-significant.

It is possible that the increased dendritic spine density observed in PirB−/− motor cortex is accompanied by an increase in the number of functional excitatory synapses. To confirm this possibility, whole-cell recordings were made from identified YFP+ L5PNs in acute coronal M1 brain slices prepared from Thy1-YFP+;PirB+/+ or Thy1-YFP+;PirB−/− mice (Figure 2A,B). A significant increase is present in the mean frequency of mEPSCs in PirB−/− mice (Figure 2C) (decreased inter-event interval, Figure 2D) but not in mean mEPSC amplitude (Figures 2E,F). These results imply that there is an increase in the number of functional excitatory inputs onto L5PNs in M1, consistent with the increase in spine density observed above. In addition, there is a significant increase in mIPSC frequency (but not mIPSC amplitude) recorded from M1 L5PNs in PirB−/− mice (Figure S2), suggesting that despite a greater number of excitatory inputs, the overall balance of excitation and inhibition is maintained in these neurons.

**Figure 2.**
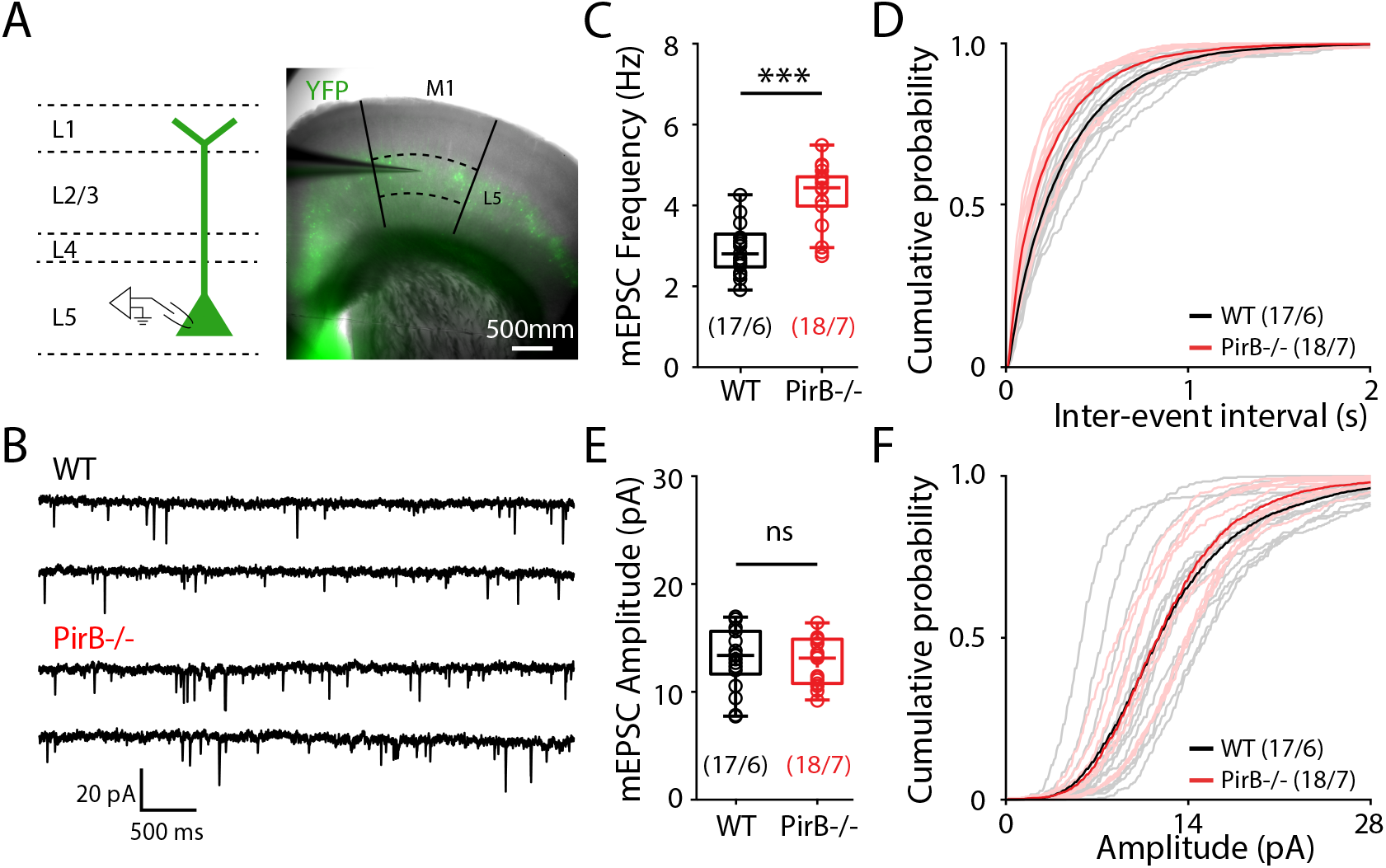
Increased Numbers of Excitatory Inputs on Layer V Pyramidal Neurons in Primary Motor Cortex of PirB−/− Mice. (A) Acute coronal slices prepared from 2-month old WT or PirB−/− mice crossed to the Thy1-YFP-H+ line. YFP expression allowed for targeting of layer 5 pyramidal neurons for whole-cell patch clamp recordings. (B) Representative recordings of mEPSCs from M1 layer 5 pyramidal neurons WT and PirB−/− mice. Tetrodotoxin (1μM) and picrotoxin (100μM) were included in the ACSF. (C, D) The frequency of mEPSCs was significantly increased (decreased inter-event interval) in PirB−/− mice, compared to WT littermates (WT: 3.151 ± 0.148; n = 17 cells / 6 mice; PirB−/−: 4.262 ± 0.179; n = 18 cells / 7 mice; p = 0.0003, Mann-Whitney). (E, F) mEPSC amplitudes were not significantly different between WT and PirB−/− mice (WT: 13.050 ± 0.671; n = 17 cells / 6 mice; PirB−/−: 12.737 ± 0.484; n = 18 cells / 7 mice; p = 0.5860, Mann-Whitney). (C-F) Circles represent individual mice. Box plots and mean ± s.e.m. are shown. *** p < 0.001; ns: non-significant.

### Enhanced Motor Learning and Motor Performance in PirB−/− Mice

To determine if increased spine density and changes in spine dynamics in M1 impact the ability of mice to learn a new motor skill, adult (2 month old) WT and PirB−/− mice were trained to perform a reaching task (Figures 3A,B and S3A) (Greenough et al., 1985; Rioult-Pedotti et al., 2000; Xu et al., 2009, Guo et al., 2015). During training, both WT and PirB−/− mice improved their reaching success rates (see Methods) in the initial 4 days (Figure 3C). However, throughout training, PirB−/− mice exhibited significantly higher rates of successful reaches compared to WT (Figure 3C). The same mice were then housed in their home cages for an additional 30 days following the training period and their performance on the same reaching task was re-evaluated. Although both pretrained WT and PirB−/− mice maintained their skillful performance, the enhanced success rate of PirB−/− mice persisted up to day 38 (Figure 3C, right). Even on day 1, within the first training session of the task, PirB−/− mice learned the task faster than the WT littermates (Figure 3D). Note that on day 1, novice WT and PirB−/− mice have the same low initial success rate (~20%) in the first 5 reaches; however, during subsequent reaches PirB−/− mice gradually outperformed their WT littermates. Consistent with a higher success rate, PirB−/− mice also completed the task significantly faster than WT throughout the 8-d training period, as well as during the re-evaluation on day 38 (Figure 3E). Moreover, a detailed analysis of reach trajectory kinematics using convolutional neural networks (DeepLabCut, see Methods) revealed that PirB−/− mice were faster to develop stereotyped, low-variability reaches compared to WT littermates (Figures 3F,G), demonstrating enhanced fine-motor adaptation in PirB−/− mice.

**Figure 3.**
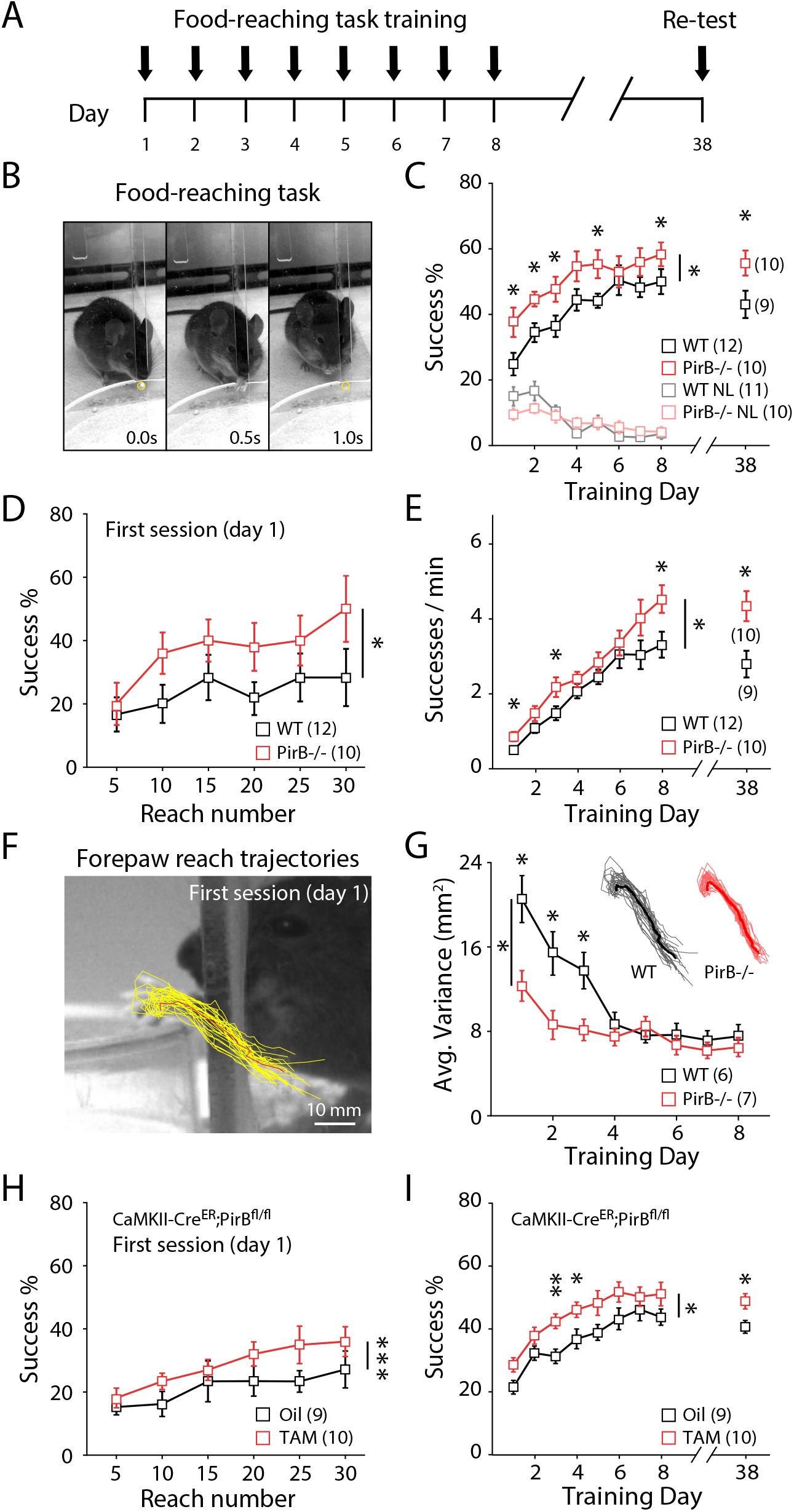
Improved Acquisition of Motor Skills in PirB−/− Mice. (A) Task timeline. 8 consecutive days of training followed by a single day of retesting 1 month later. Training begins at P60. (B) Images depicting before, during, and after a single trial of a mouse performing the reaching task. Yellow circle depicts the position of the food pellet. (C) Training days 1-8 followed by retesting at day 38. PirB−/− mice performed the reaching task better compared to WT during 8 training days (WT: n = 12; PirB−/−: n = 10; p = 0.018, 2-way repeated measures ANOVA) and again when re-assessed 30 days after training (WT: 43.056 ± 4.153%; n = 10; PirB−/−: 55.667 ± 3.818%; n = 9; p = 0.034, Mann-Whitney). Nonlearner mice (NL) are shown for both genotypes. (D) Training day 1 reach success rates, where each 5 consecutive reaches were binned. PirB−/− mice achieved a higher % rate than WT littermates within the first day of training (WT: n = 12; PirB−/−: n = 10; p = 0.029, 2-way repeated measures ANOVA). (E) PirB−/− mice made more successful reaches per minute than WT throughout 8 days of training (WT: n = 12; PirB−/−: n = 10; p = 0.037, 2-way repeated measures ANOVA) and when re-assessed 30 days later (WT: 2.794 ± 0.357; n = 10; PirB−/−: 4.343 ± 0.401; n = 9; p = 0.025, Mann-Whitney). (F) Reach trajectory kinematics analysis. Yellow: 30 individual reach trajectories. Red: average reach trajectory. (G) PirB−/− mice achieve low-variance, stereotyped reaches earlier than WT controls (WT: n = 6; PirB−/−: n = 7; p = 0.019, 2-way repeated measures ANOVA). (H) Cre-driven excision of PirB from forebrain pyramidal neurons. At P42, CaMKII-CreER;PirB^fl/fl^ mice received 5 consecutive days of tamoxifen (4 mg / day, or oil vehicle control). At P60, mice were trained and performance was measured, as above. PirB excision at older ages resulted in increased success % within first day of reaching task training (Oil: n = 9; Tamoxifen: n = 10; p = 0.0008, 2-way repeated measures ANOVA). (I) PirB excision from forebrain pyramidal neurons at older ages resulted in enhanced motor learning throughout 8 days of training (oil: n = 9; tamoxifen: n = 10; p = 0.022, 2-way repeated measures ANOVA) and when re-assessed 30 days later (p = 0.029, Mann-Whitney). (C-G) Squares depict mean (± s.e.m.) within each genotype. * p < 0.05, ** p < 0.01, *** p < 0.001.

To examine if deletion of PirB specifically from forebrain pyramidal neurons is sufficient to endow mice with enhanced motor learning, a neuronal specific deletion of PirB was generated by crossing PirB^fl/fl^ mice with a CaMKIIa-Cre^ER^ line (Madisen et al., 2009), allowing for cell-type-specific excision of PirB in the pyramidal neurons upon administration of tamoxifen at P42. 2-month-old tamoxifen- or oil-treated control CaMKIIa-Cre^ER^;PirB^fl/fl^ mice were then trained on the reaching task (Figures S3B,C). Tamoxifen-treated, PirB conditional deletion mice achieved a significantly higher success rate within the first training day compared to oil-treated controls (Figure 3H). Furthermore, conditional PirB deletion was sufficient to enhance motor learning throughout the 8-day training period and at re-evaluation 30-days later (Figure 3I), similar to germline knockout of PirB. These results demonstrate that specific deletion of PirB from forebrain pyramidal neurons is sufficient to enhance motor performance in this reaching task. In addition, because tamoxifen administration and behavioral training were performed in adult mice, these results reveal an ongoing repressive function of PirB on motor performance and they also rule out the possibility of early developmental effects and/or potential compensations caused by loss of PirB in the germline.

### Survival of Learning-Induced Spines Correlates Best with Motor Performance

Previous studies have shown that dynamic changes in dendritic spines occur during motor learning (Xu et al., 2009; Peters et al., 2014; Yang et al., 2014; Roth et al. 2020). To investigate the relationship between increased spine formation, spine survival, and enhanced motor learning in PirB−/− mice, a longitudinal experiment that combined chronic *in vivo* two-photon imaging of dendritic spines with motor skill training was performed (Figure 4A). Throughout the training paradigm, the same dendritic segments of L5PNs were repeatedly imaged through a cranial window to monitor transient and long-term spine changes (Figures 4B and S4A). In WT mice that successfully learned to perform the reaching task, L5PNs in M1 contralateral to the reaching forelimb underwent a significant increase in spine formation during the early learning period (Figure 4C), followed by a significant increase in spine elimination (Figure 4D), as compared to mice undergoing the same training but were unable to learn the task which showed stable levels of spine formation and elimination (Figures 4C,D: non-learners, WT NL). These data are consistent with many previous reports showing that motor skill learning induces rapid dendritic spine formation followed by delayed and selective spine pruning in M1 L5PNs (Xu et al., 2009; Yang et al., 2009; Wang et al., 2010; Guo et al., 2015). Trained PirB−/− mice that learned the same motor skill task also underwent a significant increase in spine formation (Figure 4F). Notably however, in PirB−/− mice, this increase in formation was not accompanied by a significant increase in spine elimination (Figure 4G). Direct comparison of spine formation and elimination in WT and PirB−/− mice that learned the task revealed that motor learning induces similar increases in spine formation despite the fact that baseline spine formation is higher in PirB−/− mice (Figure 4I). Remarkably, what differs between the genotypes is that learning-induced spine elimination is significantly lower in PirB−/− learner mice compared to WT learners (Figure 4J).

**Figure 4.**
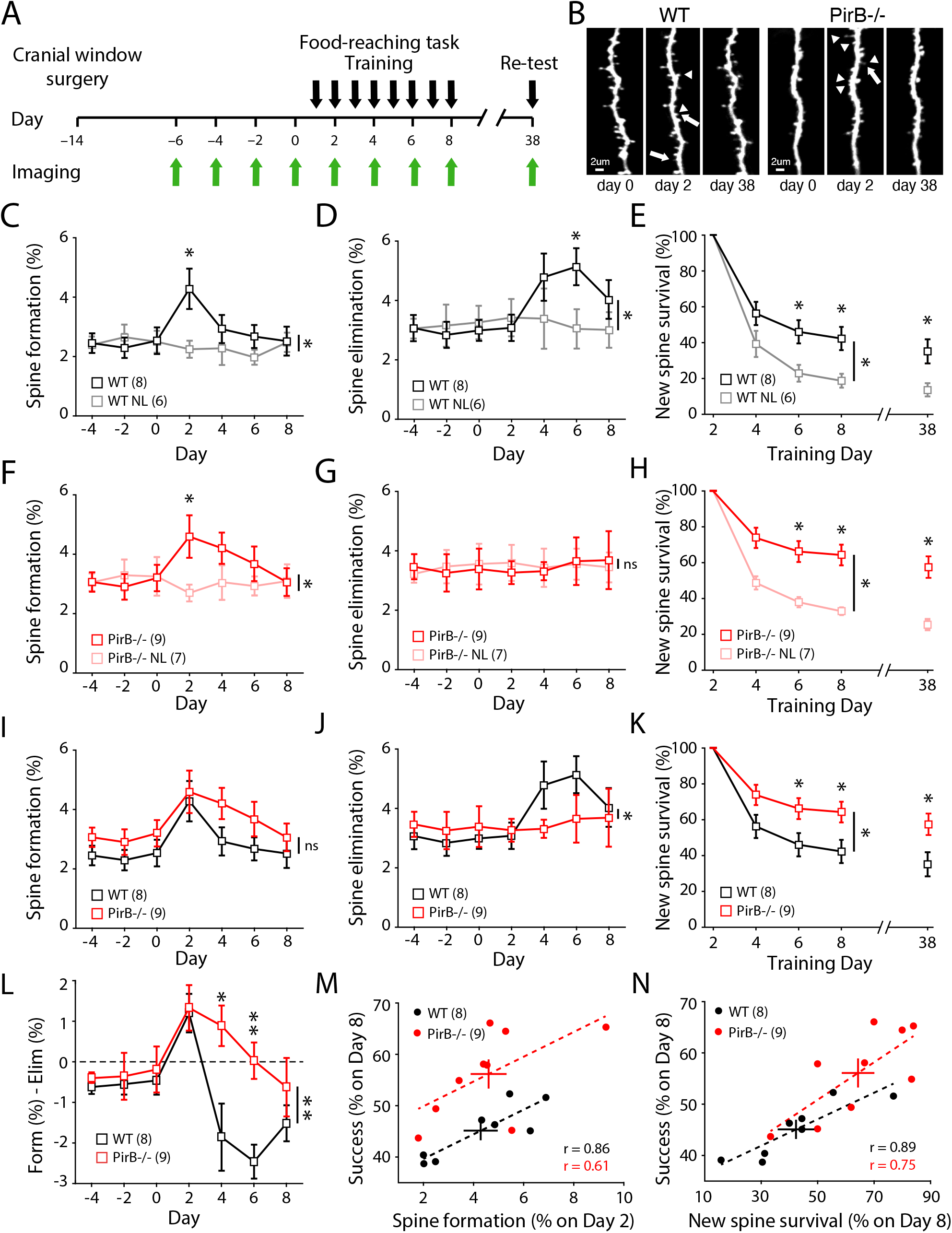
Newly Formed Spines During Early Learning are More Stable in PirB−/− Mice and the Degree of New Spine Stability Correlates with Learning Performance. (A) Experimental timeline. After recovery from cranial window implantation, mice were imaged every 2 days throughout days of pre-training, training, and 30-day post-training. (B) Representative dendrites repeatedly imaged throughout motor learning. Arrows depict spines to be eliminated, arrow heads depict newly formed spines. (C) Training resulted in a significant increase in spine formation rate in WT mice that learned the reaching task (day 0: 2.54 ± 0.45%; day 2: 4.28 ± 0.68%; n = 8; p = 0.016, Wilcoxon), which was not observed in WT mice that failed to learn (day 0: 2.51 ± 0.37%; day 2: 2.26 ± 0.29%; n = 6; p = 0.688, Wilcoxon), resulting in a significant difference in spine formation rates between WT learners and WT non-learners during training (p = 0.0084, 2-way repeated measures ANOVA). (D) Training resulted in a significant increase in spine elimination rate in WT learners (day 0: 2.99 ± 0.35%; day 6: 5.13 ± 0.62%; p = 0.008, Wilcoxon), which was not observed in WT non-learners (day 0: 3.26 ± 0.56%; day 6: 3.05 ± 0.66%; p = 0.438, Wilcoxon), resulting in a significant difference in spine elimination rates between WT learners and WT non-learners during training (p = 0.0144, 2-way repeated measures ANOVA). (E) Survival of learning-induced spines, the spines that were newly formed on day 2 (after 2 training sessions), quantified as the % of these spines persisting throughout subsequent imaging days, revealed that learning stabilized newly formed spines (p = 0.0283, 2-way repeated measures ANOVA). 30 days after the last training day, the survival of these spines was still increased in WT learners (WT learners: 35.143 ± 6.735%; WT non-learners: 13.667 ± 3.685%; p = 0.0221, Mann-Whitney). (F) Training resulted in a significant increase in spine formation rate in PirB−/− learners (day 0: 3.20 ± 0.45%; day 2: 4.60 ± 0.72%; n = 9; p = 0.008, Wilcoxon), but not in PirB−/− non-learners (day 0: 3.26 ± 0.65%; day 2: 2.70 ± 0.28%; n = 7; p = 0.375, Wilcoxon), resulting in a significant difference in spine formation rates between PirB−/− learners and PirB−/− non-learners during training (p = 0.0063, 2-way repeated measures ANOVA). (G) Training did not result in a significant increase in spine elimination rate in PirB−/− learners (day 0: 3.38 ± 0.69%; day 6: 3.65 ± 0.80%; n = 9; p = 0.734, Wilcoxon) or PirB−/− non-learners (day 0: 3.56 ± 0.69%; day 2: 3.56 ± 0.52%; p = 0.813, Wilcoxon), and spine elimination throughout training was not different between PirB−/− learners and PirB−/− non-learners (p = 0.952, 2-way repeated measures ANOVA). (H) Learning stabilized newly formed spines in PirB−/− mice (PirB−/− learners: n = 9; PirB−/− non-learners: n = 7; p = 0.001, 2-way repeated measures ANOVA). 30 days after the last training day, the survival of these spines was still increased in PirB−/− learners (PirB−/− learners: 57.556 ± 6.005%; PirB−/− non-learners: 25.333 ± 2.894%; p = 0.0028, Mann-Whitney). (I) Learning-induced spine formation rates in PirB−/− learners were not significantly different from WT learners (p = 0.228, 2-way repeated measures ANOVA). (J) Learning-induced spine elimination rates were impaired in PirB−/− learners (p = 0.035, 2-way repeated measures ANOVA). (K) PirB−/− learners have increased survival of learning-induced spines relative to WT learners (p = 0.0333, 2-way repeated measures ANOVA). 30 days after the last training day, the survival of these spines was still increased in PirB−/− learners (p = 0.033, Mann-Whitney). (L) Net dendritic spine dynamics were significantly different between WT and PirB−/− mice (WT: n = 8; PirB−/−: n = 9; p = 0.007, 2-way repeated measures ANOVA). (M) Correlation between early learning spine formation rates and last day performance on reaching task (WT: r = 0.8637; n = 8; p = 0.0057, Pearson correlation; PirB−/−: r = 0.6071; n = 9; p = 0.0830, Pearson correlation). (N) Correlation between survival of learning-induced spines and last day performance on reaching task (WT: r = 0.8900; n = 8; p = 0.0031, Pearson correlation; PirB−/−: r = 0.7455; n = 9; p = 0.0211, Pearson correlation). (C-K) Squares depict mean (± s.e.m.) within each genotype. * p < 0.05; ns: non-significant. (L-N) Circles depict individual animals, crosses depict mean ± s.e.m. of each genotype, dotted lines depict linear fit of each correlation.

It is known that newly formed task-specific dendritic spines are stabilized during training and have a significantly higher chance of survival even after motor skill training ends (Xu et al., 2009; Yang et al., 2009), forming a structural substrate that may underlie long-lasting motor memory (Hayashi-Takagi et al., 2015; Frank et al., 2018). To investigate if spine stabilization might contribute to the superior motor memory observed in PirB−/− mice, the fate of newly formed spines during early motor training was analyzed. Consistent with previous studies, in WT mice, newly-formed learning-induced spines are more stable in expert mice compared to non-learners (Figure 4E). Expert PirB−/− mice also have a higher spine survival compared to non-learners (Figure 4H). However, the percentage of newly formed spines that survive in expert PirB−/− mice is significantly greater compared with that of expert WT mice not only during the training period but also at re-evaluation one-month later (Figure 4K). When the net effect of spine turnover (spine formation - spine elimination) is compared between WT and PirB−/− mice, a rather remarkable difference emerges: spine dynamics during learning in PirB−/− mice are significantly shifted towards formation and away from elimination (Figure 4L). The net result of diminished spine elimination plus enhanced spine survival in PirB−/− mice during learning is a significant increase in spine density following training, as compared to WT (Figure S4B). These results show that learning-induced stabilization of newly formed spines is significantly enhanced in PirB−/− mice, suggesting that enhanced spine stability can serve as a potential mechanism for the improved motor learning.

Previous studies have shown that spine formation, spine elimination, and stabilization of newly formed spines, all correlate with motor learning and motor memory (Xu et al., 2009; Yang et al., 2009; Hayashi-Takagi et al., 2015). Here in our study, we have an example where both spine dynamics and learning are altered in PirB−/− mice (specifically, increased survival of learning-induced spines and enhanced motor learning). Therefore, by comparing the relationship between spine dynamics and motor learning across individual animals, it is possible to discern which factors might be the strongest predictors of motor performance. In accordance with previous studies, peak motor learning performance (% success on training day 8) correlates significantly with early-training (day 2) spine formation rates in both genotypes (Figure 4M: r = 0.86 WT; 0.61 PirB−/−). However, because learning-induced spine formation is not different between the two genotypes, it is unlikely to account for the superior performance present in PirB-/ mice. Neither pre-training spine density nor pre-training spine formation rates correlate significantly with motor learning performance in either WT or PirB−/− mice (Figures S4Cand S4D). In WT mice, learning-induced spine elimination (day 6) is significantly correlated with learning success (day 8) as previously reported, but in PirB−/− mice there is no correlation between spine elimination and motor learning (Figure S4E: r = 0.73 WT; 0.02 PirB−/−). Thus, this variable also cannot account for the superior motor learning exhibited by PirB−/− mice. Instead, by far the strongest predictor of motor learning ability is the survival of spines formed during early-training (Figure 4N: r = 0.89 WT; 0.75 PirB−/−). Indeed, the survival of these spines significantly correlates with motor performance even 30 days after the end of motor training (Figure S4F: r = 0.87 WT; 0.69 PirB−/−), and spine survival is the strongest predictor even when motor performance is measured as the variability of reach trajectory kinematics (Figures S4G-I). Taken together, these observations point to the increased stabilization of newly formed spines as the most likely explanation for enhanced motor learning in PirB−/− mice. Furthermore, they suggest that the survival of task-specific spines is the greatest general indicator of motor learning because it is not only the strongest correlation observed, but it is consistent across genotypes.

### Increased LTP and Deficient LTD in M1 L5 Pyramidal Neurons of PirB−/− Mice

Structural plasticity has been shown to depend on Hebbian synaptic plasticity (Dudek and Bear, 1993; Engert and Bonhoeffer, 1999; Matsuzaki et al., 2004; Nagerl et al., 2004; Zhou et al., 2004; Harvey and Svoboda, 2007; Bastrikova et al., 2008; Murakoshi and Yasuda, 2012). Conversely, pharmacological manipulation of long-term potentiation (LTP) and long-term depression (LTD) via targeting key signaling pathways such as Ca2+, NMDA, and dopamine, results in significant changes to dendritic spine formation, elimination, and stability (Zuo et al., 2005; Guo et al., 2015). In particular, LTP is thought to be critical for dendritic spine stabilization (Hill and Zito, 2013; Guo et al., 2015), and LTP stimulates spine formation, whereas LTD promotes spine elimination (Dudek and Bear, 1993; Engert and Bonhoeffer, 1999; Harvey and Svoboda, 2007; Bastrikova et al., 2008; Steiner et al., 2008; Kwon and Sabatini, 2011; Wiegert and Oetner, 2013). Our observations above that spine formation at baseline and stabilization following motor learning are enhanced in L5PNs in PirB−/− M1 cortex suggest that there might be alterations in Hebbian synaptic plasticity.

To determine if cellular mechanisms of synaptic plasticity are altered, coronal slices of M1 from 2-month-old Thy1-YFP;PirB+/+ or Thy1-YFP;PirB−/− mice were prepared and whole-cell recordings from identified YFP+ M1 L5PNs were made. A stimulating electrode was placed at the boundary between L1 and L2/3 of cortex and minimal stimulation was used to activate superficial layer synaptic inputs onto L5PNs (Figure 5A). Using an LTP induction protocol in which presynaptic stimulation is paired with postsynaptic depolarization (Guo et al., 2015), LTP could be reliably induced in L5PNs from WT M1 (Figure 5A), as expected. The same induction protocol in slices from PirB−/− M1 generated significantly larger LTP (Figures 5B, 5I). In both genotypes, LTP could be completely blocked by the NMDA receptor blocker 3-((*R*)-2-carboxypiperazin-4-yl)propyl-1-phosphonic acid (R-CPP, 10 μM) (Figure 5C, D, I).

**Figure 5.**
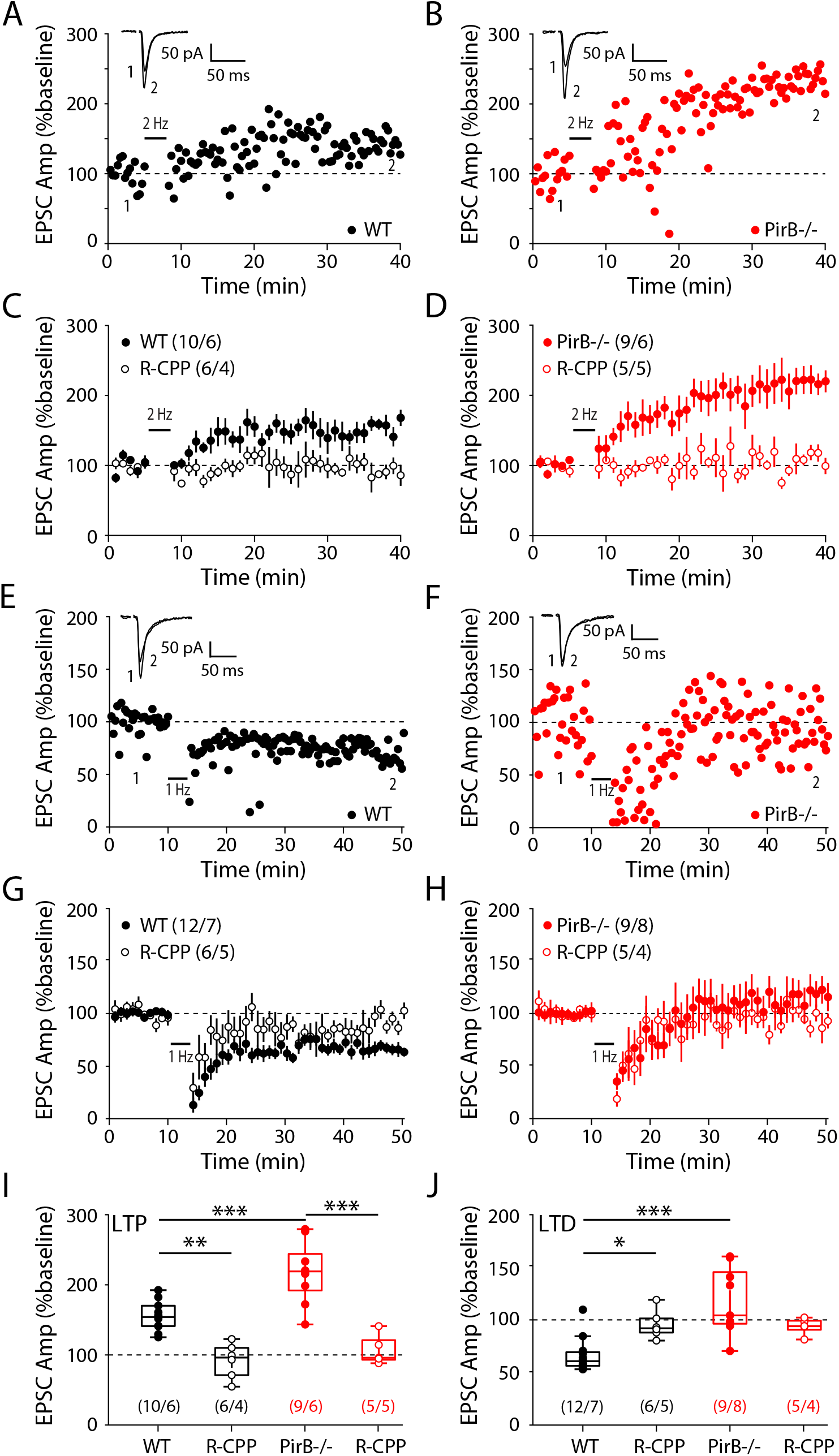
Hebbian Synaptic Plasticity Rules are Shifted in PirB−/− Mice Towards Synaptic Potentiation. (A, B) Representative whole-cell patch clamp recordings of layer 5 pyramidal neurons from acute coronal slices from WT (A) and PirB−/− (B) mice depicting evoked EPSC amplitudes before and after 2 Hz LTP induction. (C, D) Summary data showing average evoked EPSC amplitudes before and after LTP induction in WT and PirB−/− mice. A subset of recordings were performed in the presence of NMDAR antagonist R-CPP (10μM). (E, F) Representative recordings from WT (E) and PirB−/− (F) mice depicting evoked EPSC amplitudes before and after 1 Hz LTD induction. (G, H) Summary data showing average evoked EPSC amplitudes before and after LTD induction in WT and PirB−/− mice. A subset of recordings were performed in the presence of NMDAR antagonist R-CPP (10μM). (I) LTP was significantly increased in PirB−/− mice (WT: 155.127 ± 6.946%; n = 10 cells / 6 mice; PirB−/−: 217.592 ± 14.639%; n = 9 cells / 6 mice; p = 8.777e-4, 1-way ANOVA multiple comparisons). Observed LTP was NMDAR-dependent for both WT (WT: 155.127 ± 6.946%; n = 10 cells / 6 mice; WT + R-CPP: 91.849 ± 10.238%; n = 6 cells / 4 mice; p = 0.0027, 1-way ANOVA multiple comparisons) and PirB−/− mice (PirBKO: 217.592 ± 14.639%; n = 9 cells / 6 mice; PirB−/− + R-CPP: 107.266 ± 9.622%; n = 5 cells / 5 mice; p = 4.950e-6, 1-way ANOVA multiple comparisons). (J) LTD was significantly impaired in PirB−/− mice (WT: 66.345 ± 4.686%; n = 12 cells / 7 mice; PirB−/−: 117.504 ± 10.581%; n = 9 cells / 8 mice; p = 3.245e-5, 1-way ANOVA multiple comparisons). Observed LTD was NMDAR-dependent for WT (WT: 66.345 ± 4.686%; n = 12 cells / 7 mice; WT + R-CPP: 95.462 ± 5.490%; n = 6 cells / 5 mice; p = 0.043, 1-way ANOVA multiple comparisons). (A, B, E, F) Average traces are shown for baseline (black) and end of recording (red), where each is the average of the last 5 minutes of their period (15 traces). (I, J) Circles represent individual cells. Box plots depict mean (± s.e.m.) within each genotype. * p < 0.05, ** p < 0.01, *** p < 0.001.

Next, LTD was assessed. Coronal brain slices of M1 from 1 month old mice were used because previous studies have indicated that LTD in L5PNs cannot be elicited in older mice using a pairing protocol (Guo et al., 2015). In our hands, this low-frequency (1 Hz) pairing LTD protocol successfully induced NMDA receptor dependent LTD in WT mice (Figure 5E and 5G). However, the same protocol failed to induce LTD in PirB−/− mice (Figures 5F, H, J).

These observations demonstrate that L5PNs in PirB−/− mice have altered Hebbian synaptic plasticity rules in M1 that favor synaptic potentiation over synaptic depression, a result consistent with previous observations of changes in Hebbian synaptic plasticity present in PirB−/− in visual cortex (Djurisic et al., 2013) and hippocampus (Djurisic et al., 2019), structures in which pyramidal neurons also normally express PirB. These results also demonstrate that one function of PirB is to regulate cellular mechanisms of synaptic plasticity, which ultimately may account for the increases in stabilization of learning-induced spines and the enhancement in motor learning observed in PirB−/− mice.

### Enhanced Motor Learning and Spine Survival with Acute Blockade of PirB Function in M1 of Wild Type Mice

The improved motor learning along with increased survival of learning-induced spines in PirB−/− mice suggests that PirB normally acts to limit motor learning. This consideration made us wonder if it might be possible to lift limits in adult WT mice by acutely blocking PirB. To accomplish this goal, a classic protein-based approach was used in which a recombinant soluble PirB decoy receptor (sPirB) was generated (Bochner et al., 2014). This decoy receptor consists of only the PirB ectodomain (containing all six immunoglobulin (Ig)-like domains) plus Myc- and His-tags for purification and detection. sPirB functions by binding all endogenous PirB ligands, thereby blocking activation of full length PirB downstream signaling. “Decoy receptors” have been used very effectively as drugs in human clinical application (Davis et al., 1994; Cabelli et al., 1997; Holash et al., 2002), and previous work has demonstrated that a 2 week infusion of sPirB into visual cortex of adult WT mice effectively blocks endogenous PirB proximal signaling *in vivo*, reinstating juvenile-like ocular dominance plasticity and facilitating recovery from Amblyopia (Bochner et al., 2014).

To examine if this pharmacological blockade of PirB can alter spine dynamics and improve motor learning, osmotic minipumps were implanted in 2 month old Thy1-YFP-H mice and sPirB or BSA control (a similar sized protein to sPirB) was infused into M1 cortex contralateral to the imaged forelimb over a period of 2 weeks. As above, chronic two-photon imaging was then used over an additional 8 day infusion period to assess the effect of acute blockade of PirB on dendritic spine dynamics and motor learning (Figure 6A). At the completion of the experiment, in some animals the brain was removed, sectioned, and immunostained using an anti-Myc antibody to identify the cortical area infused with sPirB (Figure 6B). In all cases, Myc- immunostaining was restricted to a localized 3-4 mm^3^ region including M1.

**Figure 6.**
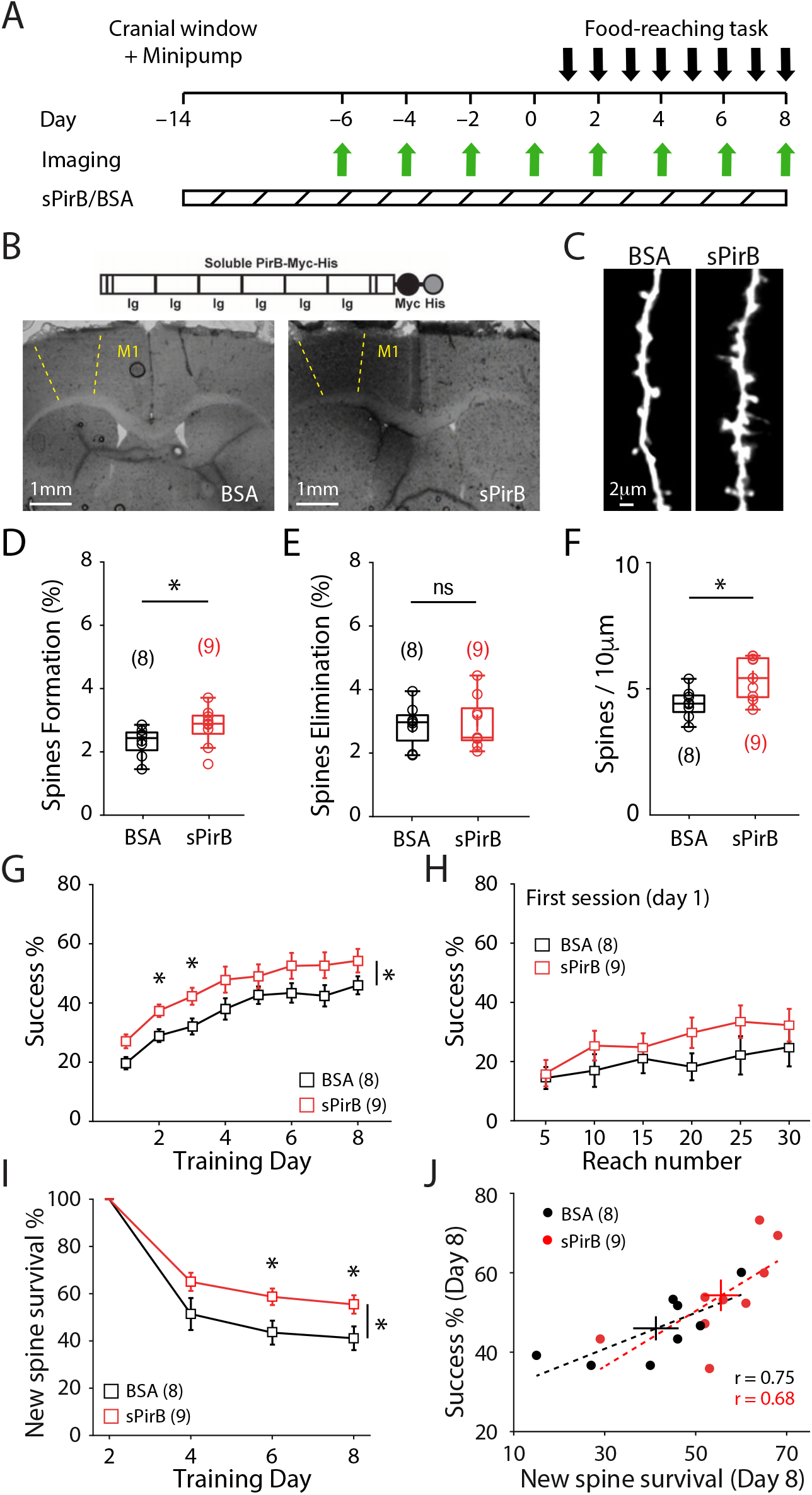
Chronic Infusion of sPirB into M1 of WT Mice Enhances Stability of Newly Formed Spines and Improves Performance on Motor Skill Training. (A) Experimental timeline. After recovery form cranial window and osmotic minipump implantation, WT mice are repeatedly imaged with 2-day intervals before and during reaching task training. (B) (Top) Schematic of sPirB structure depicting extracellular Ig domains of PirB. (Bottom) alkaline phosphatase staining of sPirB-Myc depicting positive staining in M1 of animal infused with sPirB (right) vs. a control mouse infused with BSA (left). (C) Representative dendrites imaged after 2 weeks of chronic BSA (left) or sPirB (right) infusion. (D, E) Increased baseline rate of spine formation (BSA: 2.331 ± 0.162%; n = 8 mice; sPirB: 2.820 ± 0.204%; n = 9 mice; p = 0.036, Mann-Whitney), but not spine elimination (BSA: 2.844 ± 0.238%; sPirB: 2.910 ± 0.270%; p = 0.942, Mann-Whitney) after chronic infusion of sPirB. (F) Chronic infusion of sPirB significantly increased dendritic spine density of WT mice (BSA: 4.310 ± 0.208; n = 8 mice; sPirB: 5.343 ± 0.285; n = 9 mice; p = 0.021, Mann-Whitney). (G) WT mice infused with sPirB performed the reaching task better compared to WT mice infused with BSA during 8 training days (BSA: n = 8; sPirB: n = 9; p = 0.034, 2-way repeated measures ANOVA). (H) sPirB infusion did not significantly enhance motor learning within the first training day (BSA: n = 8; sPirB: n = 9; p = 0.051, 2-way repeated measures ANOVA). (I) sPirB infusion significantly increased the survival of learning-induced spines, the spines that were newly formed on day 2 (after 2 training sessions), quantified as the % of these spines persisting throughout subsequent imaging days (BSA: n = 8; sPirB: n = 9; p = 0.0436, 2-way repeated measures ANOVA). (J) Correlation between survival of learning-induced spines and last day performance on reaching task (BSA: r = 0.7506; n = 8; p = 0.0319, Pearson correlation; sPirB: r = 0.6817; n = 9; p = 0.0431, Pearson correlation). Circles depict individual mice, crosses depict mean ± s.e.m. of each genotype, dotted lines depict linear fit of each correlation. (D, E, F) Circles represent individual mice. Box plots and mean ± s.e.m. are shown. (G, H, I) Squares depict mean (± s.e.m.) within each genotype. (D-I) * p < 0.05, ns: non-significant.

A brief two-week sPirB infusion into M1 of WT mice can increase spine formation rate (Figures 6C,D) with no detectable effect on spine elimination (Figure 6E), resulting in a significant increase in spine density (~24%; Figures 6C,F). Next, sPirB or BSA (used as a control because it is of similar size to sPirB)-infused Thy1-YFP-H mice were trained on the reaching task and apical dendritic segments of L5PNs in M1 were repeatedly imaged (Figure 6A). Remarkably, there was a significant enhancement in motor learning in mice receiving sPirB infusion compared to mice infused with BSA throughout the 8-d training period (Figure 6G). The first training session on day 1 did not reveal a significant enhancement in performance in sPirB vs BSA treated mice (Figure 6H), possibly due to the fact that sPirB infusion had only started 14 days earlier and time is needed for levels to achieve an effective blockade in M1. Nevertheless, by training day 6 a significant increase in survival of newly formed spines by training day 6 was observed in sPirB infused mice compared to BSA infused controls (Figure 6I), recapitulating our findings in germline PirB−/− mice. Consistent with those findings, a significant correlation between the magnitude of new spine survival and motor performance was also seen (Figure 6J). These observations extend our study beyond genetic manipulations of PirB to show that a relatively simple functional blockade of PirB using a protein-based approach yields similar results in adult WT mice: not only is an increase in spine survival by acute and localized blockade of PirB in M1 adult mice sufficient to enhance motor learning, but also the survival of learning-induced spines is the strongest structural correlate of motor learning.

## DISCUSSION

There are several major findings from this study. Together they provide key evidence supporting the idea that increasing the stability of newly formed spines is sufficient to enhance motor learning. By investigating PirB−/− mice, we have identified a neuronal receptor that regulates dendritic spine dynamics and motor learning. Targeting PirB genetically or pharmacologically significantly increases baseline spine formation on the apical dendrites of L5PNS in M1. Moreover, newly formed spines generated during motor skill training are more stable than normal, and motor learning itself is enhanced. Notably, the greatest predictor of motor learning across animals and genotypes is the magnitude of spine survival. At the synaptic level, Hebbian plasticity rules in M1 are shifted towards potentiation and away from depression, providing a likely mechanism for the enhanced stabilization of dendritic spines during learning in PirB−/− mice. Finally, we have used 2 different methods to demonstrate that motor learning can be enhanced by selectively manipulating PirB just in cortex. First, motor learning is enhanced in mice lacking PirB only in forebrain pyramidal neurons (Figure 3 H,I). Second, infusing sPirB to block endogenous PirB signaling within M1 of adult WT mice recapitulates both the increased spine survival and enhanced motor learning. Our work adds to a growing theory that the formation and stabilization of dendritic spines are the chief mechanisms by which motor skills are acquired and maintained (Yang et al., 2009; Peters et al., 2017). Results here also provide an alluring target – PirB – for increasing dendritic spine stability: here we have discovered that PirB blockade not only enhances motor learning in the healthy brain but also may counteract or even prevent phenotypes of spine instability and accelerated elimination observed in various neurological and psychiatric disorders.

### Stability of learning-induced spines as a strong candidate for a structural correlate of motor learning

Here we find that for each genotype, the magnitude of learning-induced spine formation predicts performance (Figure 4M). This result is consistent with previous studies revealing a significant correlation between learning-induced spine formation and motor learning (Xu et al., 2009; Yang et al., 2009). However, spine *formation* cannot predict the *enhanced* motor learning present in PirB−/− mice because learning-induced spine formation rates do not differ between WT and PirB−/− mice (Figure 4I). Instead, the *survival* of learning-induced newly formed spines stands out as the major factor correlating with motor learning: increased spine survival is sufficient to predict performance of individual animals regardless of genotype (Figure 4N). These observations therefore allow a more complete understanding of the relative importance of different parameters of spine dynamics to motor learning (Ziv and Brenner, 2018). Indeed, in WT mice, simply increasing the stability of newly formed spines by infusing sPirB is sufficient to enhance their learning in a predictable manner (Figures 6I,J).

L5PNs in PirB−/− motor cortex have significantly more dendritic spines than WT mice (Figure 1E), consistent with previous observations that spine pruning on dendrites of pyramidal neurons in visual cortex is deficient during development (Djurisic et al., 2013; Vidal et al., 2016). However, it cannot be argued that merely increasing spine density is sufficient to improve motor learning. Indeed, Fragile X mouse models also exhibit significantly increased spine numbers, yet these mice have significant learning deficits (Padmashri et al., 2013). Furthermore, a significant relationship between spine density and motor learning is not evident across the animals used in this study, regardless of genotype (Figure S4C). Instead, we find that enhanced learning in PirB−/− mice is more likely due to the increased survival of newly formed spines, a relationship that holds true across all animals of either genotype (Figure 4N). A corollary of this logic is consistent with the observation that spine instability accompanies learning deficits in Fragile X mice. Furthermore, dendritic spines in Fragile X mouse cortex are significantly thinner and smaller in size (Cruz-Martin et al., 2010), whereas the spines sizes observed in PirB−/− mice are not different from those in WT mice (Bochner et al., 2014). It is likely that the increased spine stability present in PirB−/−, combined with significantly increased spine formation rates, results in the elevated spine numbers.

### Enhanced motor learning can occur even when spine elimination is deficient

Previous studies have shown that spine elimination correlates with, and may even be required, for motor learning (Yang et al., 2009; Chen et al., 2015). Although a significant correlation between learning-induced spine elimination and motor learning is observed here in WT mice as expected (Figure S4E), such a correlation was notably absent in PirB−/− mice, despite their enhanced performance on the reaching task. It is important to note that at baseline, spine elimination on L5PNs in PirB−/− mice is comparable to that seen in WT mice. In fact, the difference appears to lie specifically in the learning-induced elimination engaged following training. Although the overall magnitude of spine elimination in PirB−/− is diminished, it remains possible that selective elimination of specific spines still occurs. Nevertheless, the spine dynamics observed here in PirB−/− mice demonstrate that superior acquisition of a motor skill can occur even in the face of diminished spine elimination.

### Altered Hebbian plasticity as a mechanism for increased spine stabilization and enhanced learning

Results from this study of L5PNs in PirB−/− mice link altered learning and spine dynamics with altered mechanisms of Hebbian synaptic plasticity. While there is growing evidence for a significant relationship between spine dynamics and learning from studies of mutant mice and from directly altering spine dynamics in WT mice (Hayashi-Takagi, et al., 2015; Frank et al., 2018), few studies have examined changes in all 3 parameters (Xu et al., 2009; Yang et al., 2009). Studies have connected altered learning to altered mechanisms of LTP and LTD (Clem et al., 2008; Zhou et al., 2016), and previous work has shown that PirB−/− mice display altered Hebbian plasticity rules that favor synaptic potentiation over synaptic depression in visual cortex and hippocampus (Djurisic et al., 2013; Djurisic et al., 2019). Therefore, one might expect that altered Hebbian plasticity rules in PirB−/− cortex would translate to altered spine dynamics favoring stabilization, which is a likely downstream readout of enhanced LTP (see below). Here we demonstrate that L5PNs in M1 of PirB−/− mice also exhibits greater LTP and impaired LTD, and further, that this change accompanies significantly increased spine stability and enhanced motor learning.

The significant increase in spine clustering observed in PirB−/− mice (Figure S1D) could also be related to the shift in Hebbian synaptic plasticity favoring LTP. Indeed, there is growing evidence that synaptic clustering can lower local thresholds for LTP (Harvey and Svoboda, 2007; Murakoshi and Yasuda, 2012). Clustering of synaptic inputs can lead to nonlinear integration of functionally similar or dissimilar inputs, resulting in greater influence on the overall information encoding of the neuron (Froemke et al., 2005; Larkum and Nevian, 2008; Wilson et al., 2016; Iacaruso et al., 2017; El-Boustani et al., 2018; Frank et al., 2018). A similar increase in spine clustering is also observed in visual cortex of PirB−/− mice using an automated spine analysis algorithm (Xiao et al., 2018; Xiao et al., unpublished). Though more is known about the role of spine clustering in sensory areas such as visual cortex (Lee et al., 2019), further studies are needed to characterize the functional consequence of clustered spines on L5PNs in M1 (Fu et al., 2012; Gokce et al., 2016).

The PirB receptor has intracellular signaling domains that interact with SHP2 (Takai, 2005; Syken et al., 2006) and also with the serine-threonine phosphatases PP2A and PP2B/calcineurin, and their target, cofilin (Kim et al., 2013). PirB downstream signaling via cofilin dephosphorylation is thought to drive postsynaptic actin filament disassembly and synapse collapse, a mechanism that is accentuated in AD mouse models and disengaged in the visual cortex and hippocampus of PirB−/− mice (Kim et al., 2013; Djurisic et al., 2019). Our observations in M1 L5PNs are consistent with this model: LTD induction protocols that are sufficient to induce robust synaptic depression in WT mice appear entirely ineffective in PirB−/− mice. This loss of LTD may account for the reduced learning-induced spine elimination and enhanced spine stabilization observed in PirB−/− mice, which then would account for faster and enhanced motor learning.

### Increased excitatory and inhibitory tone in PirB−/− L5PNs

Whole cell recordings from L5PNs reveal that the frequency of mEPSCs is increased in PirB−/− mice (Figure 2). This increase in mEPSCs is entirely consistent with the increased spine density observed here and implies that there is an increase in the number of functional excitatory inputs to L5PNs. We also observed a significant increase in the frequency of mIPSCs (Figure S2). This increase may be a result of altered inhibitory plasticity, and/or a consequence of a homeostatic upscaling of inhibitory inputs in response to the increased number of excitatory inputs to stabilize excitation and E/I balance (Froemke et al., 2007; Maffei and Turrigiano, 2008). Spine dynamics, elimination in particular, have been shown to be regulated by inhibitory inputs. GABA signaling along dendrites locally promotes LTD and spine elimination (Hayama et al., 2013) and inhibitory inputs undergo dramatic structural and functional plasticity during learning (Donato et al., 2013; Chen et al., 2015). Further investigation of inhibitory networks and dynamics in mice may elucidate additional mechanisms by which the stability of dendritic spines is enhanced in PirB−/− mice. Whatever the mechanism, E/I balance is preserved with loss of PirB function and the elevated number of functional GABAergic inputs may compensate for increased excitation to prevent circuit instability.

### Specificity of enhanced motor learning revealed by infusing sPirB into adult WT M1

Motor learning is a distributed process involving many structures and modalities (Jin and Costa, 2010; Guo et al., 2015; Makino et al., 2017; Lemke et al., 2019; Roth et al., 2020). Here we made a focal blockade of endogenous PirB restricted to M1, demonstrating that targeting this site with local infusion of sPirB in WT mice is sufficient to enhance motor learning. But it is also important to consider the identity of the presynaptic partners of dendritic spines, as there is a great heterogeneity of inputs to L5PNs that remains to be explored. Notably, previous work has shown that motor learning induces plasticity of horizontal connections within motor cortex (Rioult-Pedotti et al., 2000), as well as of thalamocortical synapses onto a subset of functionally relevant L5PNs (Biane et al., 2016). Dopaminergic inputs to M1 L5PNs have also been shown to regulate synaptic plasticity and spine dynamics, emphasizing the multitude of levels of regulation surrounding this process (Guo et al., 2015). Further characterization of these various inputs, and their own plasticity mechanisms, will provide a more complete picture of the dynamic synaptic rewiring during motor learning. Nevertheless, we show that manipulating spine stability locally in M1 is sufficient to result in significant enhancements to motor performance, even in WT adult. These results imply that intervening in one place can improve performance and may result in a cascade of changes throughout the motor system. These considerations also suggest that sPirB or other approaches that decrease PirB function within cortex can be leveraged to counteract, or even prevent, the detrimental spine instability and loss that accompanies many neurological disorders such as PD (Stephens et al., 2005; Guo et al., 2015) and AD (Tsai et al., 2004; Spires et al., 2005; Shankar et al., 2007). Learning and memory deficits are hallmarks of such pathologies involving dendritic spines loss, and it may be possible to restore or protect these functions by infusion of sPirB into the affected brain regions.

## ACKNOWLEDGMENTS

We wish to thank Dr. Maja Djurisic and Dr. Richard Roth for their invaluable advice and careful reading of the manuscript, and Nora Sotelo-Kury, Peggy Kemper, Christina Chechelski, and Yu Liu for technical support. This work was funded by grants from the NINDS/NIH NS075136 (J.B.D.), the Klingenstein Foundation (J.B.D.), NIH EY02858 (C.J.S.), the Mathers Charitable Foundation (C.J.S.), HHMI Gilliam Fellowship (E.A.), NSF GRFP (E.A.), and DARE Fellowship (E.A.)

## AUTHOR CONTRIBUTIONS

E.A., C.J.S., and J.B.D. designed experiments. E.A. performed behavioral, imaging, and electrophysiology experiments. E.A. designed analysis software and analyzed data. E.A. and A.R. performed cranial window and osmotic pump implantation surgeries. O.J. aided with acquisition of preliminary data. A.R. performed sPirB purification, and alkaline phosphatase experiments. E.A., C.J.S., and J.B.D. wrote the manuscript with input from all authors.

## LEAD CONTACT AND MATERIALS AVAILABILITY

This study did not generate new unique reagents. Further information and requests for resources and reagents should be directed to and will be fulfilled by the Lead Contacts, Carla J. Shatz and Jun B. Ding.

## EXPERIMENTAL MODEL AND SUBJECT DETAILS

### Animals

All experiments were performed in accordance with protocols approved by the Stanford University Animal Care and Use Committee in keeping with the National Institutes of Health’s *Guide for the Care and Use of Laboratory Animals*. Animals were kept at a 12hr:12hr light/dark cycle. All behavior and imaging experiments were carried out at consistent times of day (behavior: 7pm-10pm, every day; imaging: 7am-10am, every 2 days) to avoid circadian and sleep related effects. Both male and female mice were used for all experiments (P60-P80, with the exception of LTD recordings performed at P20-22). PirB−/− and PirB^fl/fl^ mice were generated as previously described (Syken et al., 2006; Djurisic et al., 2019). PirB+/− lines were maintained to generate littermate PirB+/+ and PirB−/− mice used in all experiments (P60-P80). The CaMKIIa-Cre^ER^ line was obtained from Jaxon Labs (JAX 012362; Madisen et al., 2009). CaMKIIa-Cre^ER^;PirB^fl/fl^ mice were crossed to PirB^fl/fl^ mice to yield 50% Cre+ animals.

## METHODS DETAILS

### Cranial window surgery

Chronic cranial window was implanted over M1 of P45-P50 as previously described (Holtmaat et al., 2009). Briefly, mice were anesthetized with isoflurane and anti-inflammatory drugs Carprofen and Dexamethasone were administered. The head of the animal was stabilized in a stereotaxic frame. The skin and periosteum were removed to expose the skull from the olfactory bulb to the cerebellum. A high-speed drill was used to drill a circular groove in the skull over the left primary motor cortex. Drilling was intermittent for heat dissipation and cool sterile saline was periodically added to the skull to avoid damage to the underlying cortex due to friction-induced heat. When the remaining island of cranial bone moved easily in response to light touch it was removed and a sterile circular 3mm round window was placed over the exposed cortex. Animals with compromised dura and/or vasculature were excluded. A thin layer of cyanoacrylate was applied to the surface of the skull and the edge of the cover glass to seal off the exterior, followed by dental resin to cover both the exposed skull and wound edge. Lastly, a titanium bar with threaded screw holes was attached to the skull postlateral to the cranial window for stabilizing the head during subsequent imaging sessions. Animals recovered from surgery for 2 weeks before imaging experiments.

### Single-pellet reaching task

The single-pellet reaching task for testing motor learning was conducted as previously described (Xu et al., 2009). Briefly, before training, animals (male and female, P45-P55) were food restricted to 90% of their free-feeding body weight. The training chamber was constructed using transparent Plexiglas. A vertical slit was made on the front side of the chamber for the mouse to reach through for millet seeds. The experiment involved a shaping phase followed by a training phase. In the shaping phase, mice were placed in the reaching chambers to familiarize with the new environment and task requirements. On the last day of shaping, the dominant forelimb of the animal was determined by placing a pile of seeds in front of the slit and letting the animal reach using either limb. When at least 20 reaches were conducted within 20 min, and limb preference met 70% or greater, dominant forelimb was determined. All mice finished shaping within 3-7 days. Every training day consisted of 30 trials with the dominant forelimb, or 20 min. Reaches were classified as a ‘success’ if the animal used the dominant forelimb to grasp the seed and bring it to its mouth, a ‘drop’ if the animal used the dominant forelimb to grasp the seed but dropped it when retrieving it, or a ‘fail’ if the animal used the dominant forelimb to reach for the seed but failed to grasp it. Success rates was quantified as the percentage of ‘success’ reaches over total reaches. Speed of success was quantified as the number of ‘success’ reaches divided by the total session time (20 min max) as many of the animals saturated their ‘success’ rate performance but continued to increase the speed of their ‘success’. The training phase lasted 8 days, with a final session conducted 30 days after the last training session. All shaping and training sessions were conducted at similar times of day (7pm – 10pm). Controls were ‘non-learners’: mice that underwent shaping but failed to reach a minimum of 20% success rate throughout training. Mice that underwent complications after surgery or during food restriction were excluded from the study.

### Tamoxifen administration

Tamoxifen (Sigma T5648) was dissolved at 20mg/ml in corn oil (Sigma C8267) by 60 second vortex and overnight inversion at 37°C and stored in aliquots at 4°C. 2 mg (100ul) tamoxifen (or oil vehicle) was injected intraperitoneally into P42 CaMKIIa-Cre^ER^;PirB^fl/fl^ mice at 8am and again at 8pm daily (4 mg / day), for 5 consecutive days, with the last injection being ≥ 2wks before behavioral training, at which point the mice were 2-months old.

### Analysis of reaching kinematics

Videos of reaching sessions were recorded at 100 Hz using a Grasshopper3 camera (FLIR Systems) and chopped to include only reaching bouts (150-300 frames each). X-Y paw positions were then extracted using DeepLab Cut (Mathis et al., 2018) using a Linux workstation utilizing an NVIDIA GeForce RTX 2080TI graphics card. Briefly, the DeepLab Cut model was trained using ~500 manually annotated frames, after which the model output was checked for errors and corrected. After X-Y data was extracted for each bout, trajectory variance was quantified as the average of the X and Y trajectory variances, which were calculated by taking the average pairwise variances of reaches in the X and Y dimensions across all reaches for that training session (~30 reaches / day).

### In vivo two-photon imaging

Chronic two-photon imaging experiments were performed as previously described (Xu et al., 2009; Holtmaat et al., 2009; Guo et al., 2015). Repeated imaging of apical dendritic stretches of L5 pyramidal neurons 10-100μm below the cortical surface was performed through the cranial window using a custom-built two-photon microscope or an Olympus microscope (FV1200) with a Mai Tai Ti:sapphire laser (Spectra-Physics) at 920 nm with a low laser power (output optical power <40 mW) to avoid phototoxicity. Image stacks were taken with a z-step of 0.7 μm using a water-immersion Olympus objective. In order to re-locate the same region across imaging days, lower magnification dendritic structures of interest were taken with a 2.0 μm step size and brain vasculature of the area was imaged with a CCD camera to serve as landmark guides. After imaging sessions, the titanium bar was detached from the metal base and the animal was returned to the original home cage. Imaged regions were determined using stereotaxic coordinates (Tennant et al., 2011) (1.2 mm lateral from the midline, 1.3 mm anterior to the bregma).

### Identification and presentation of dendritic spines

As previously described (Xu et al., 2009; Guo et al., 2015), individual dendritic protrusions (minimum length of 1/3 the dendrite diameter) were manually identified and tracked along dendritic segments. Spine changes (i.e. formation and elimination) were determined by comparing two temporally adjacent images of the same spines. Spines were classified as stable if they were present in both images, using their spatial relationship to landmarks and/or their position relative to other spines. An eliminated spine was one that was present in the first image but not detected on the second image. A newly formed spine was one that was absent in the first image but appeared in the second image. Representative dendrites were created by making 2D projections of 3D stacks containing in-focus dendritic segments. Projections were thresholded, contrast-enhanced, and median filtered to create the final images.

### Dendritic spine clustering analysis

Spine clustering was analyzed by quantifying nearest neighbor (NN) distance: for each spine, the distance along the dendrite to the nearest adjacent spine (Fu et al., 2012; Gokce et al., 2016). Expected NN distributions were generated by simulating dendrites with shuffled spine positions and spine densities matching the observed values from imaging experiment, and extracting the corresponding NN values. Repeating this process through 1000 simulations generated expected NN distributions for WT and PirB−/− mice given their respective spine densities (Figure S1D, shaded). Spine clustering was then quantified as significant differences between the observed NN distributions and the expected NN distributions for each genotype.

### Whole-cell slice electrophysiology

Mice (male and female, 2 months old) were anesthetized with isoflurane, decapitated, and brains were extracted and briefly submerged into chilled artificial cerebrospinal fluid (ACSF) containing 125 mM NaCl, 2.5 mM KCl, 1.25 mM NaH2PO4, 25 mM NaHCO3, 15 mM glucose, 2 mM CaCl2, and 1 mM MgCl2, oxygenated with 95% O2 and 5% CO2 (300-305 mOsm, pH 7.4). Coronal slices (300 μm thickness) containing M1 were then prepared using a tissue vibratome (VT1200S, Leica), incubated in chambers containing 34°C ACSF for 30 min, and then allowed to recover at room temperature for 30 min. After recovery, slices were transferred to a submerged recording chamber perfused with ACSF at a rate of 2-3 ml/min at a temperature of 30-31°C. Motor cortex M1 layer V pyramidal neurons were identified by YFP fluorescence (BX51, Olympus). Whole-cell voltage clamp recordings were made with glass pipettes (3-4 MΩ) filled with internal solution containing 126 mM CsMeSO_3_, 10 mM HEPES, 1 mM EGTA, 2 mM QX-314 chloride, 0.1 mM CaCl_2_, 4 mM Mg-ATP, 0.3 mM Na_3_-GTP, and 8 mM disodium phosphocreatine (280-290 mOsm, pH 7.3 with CsOH), and cells were voltage clamped at −70 mV. Access resistance was 15-25 MΩ for all recordings and only cells with a change in access resistance <20% throughout the entire experiment were included in the analysis. Miniature excitatory postsynaptic currents (mEPSCs) were measured by continuously recording for 10 min in the presence of 1 μM Tetrodotoxin to prevent action potential firing and 50 μM Picrotoxin to block GABAA receptor-mediated currents. For miniature inhibitory postsynaptic currents (mIPSCs), 10 μM NBQX and 10 μM (R)-CPP were added to block glutamate receptors and NMDA receptors, respectively. For pairing LTP, induction stimuli were delivered within 13 min of achieving whole-cell configuration. After a stable baseline (5 min), presynaptic stimulation (2 Hz, 360 pulses) was paired with postsynaptic depolarization to +10 mV. For pairing LTD (3-week old mice), induction stimuli were delivered within 20 min of achieving whole-cell configuration. After establishing a stable baseline (10 min), low-frequency stimulation (1 Hz, 200 pulses) was paired with postsynaptic depolarization to −30 mV. EPSCs were induced by stimulating the superficial layer of M1 via a concentric bipolar stimulating electrode (FHC) located at the border between L1 and L2/3 (Guo et al., 2015). EPSCs were evoked at 0.05 Hz and three successive EPSCs were averaged and quantified relative to the normalized baseline. Tissue surrounding the recording area were cut to prevent polysynaptic responses. Picrotoxin (50 μM) was present in the ACSF perfusion for LTP and LTD experiments. Whole-cell patch clamp recordings were performed using a Multiclamp 700B (Molecular Devices), monitored with WinWCP (Strathclyde Electrophysiology Software) and analyzed offline using Clampfit 10.0 (Molecular Devices) and custom-made MATLAB (Mathworks) software. Signals were filtered at 2 kHz and digitized at 10 kHz (NI PCIe-6259, National Instruments).

### Chronic minipump infusion of sPirB

For sPirB infusion experiments (Figures 6, S6), osmotic minipumps (Alzet, 0.2ul/hr, 100ul capacity) were implanted as described previously (Bochner et al., 2014) with cranial windows in the same surgery. Approximately 16-18 hours before cranial window surgeries, minipumps were filled with either sPirB (1mg/ml) or BSA (1mg/ml) (VWR, EM-2930) in 1x PBS and incubated at 37°C overnight. The next day, minipumps were implanted subcutaneously and connected to a cannula (PlasticsOne) which was inserted directly posterior to the cranial window. A thin layer of cryanoacrylate glue and dental cement resin was used to secure cannulas to stereotaxic frame for the duration of imaging sessions and behavioral testing.

### Immunohistochemistry of sPirB

At the completion of live imaging experiments, mice were euthanized and whole brains were rapidly frozen in M-1 embedding matrix. Coronal sections were cut at a thickness of 15μm using a cryostat and mounted onto slides for processing. To stain for sPirB-myc, slices were fixed and stained as performed previously (Bochner et al., 2014). Briefly, slices were submerged in 4% PFA and fixed for 30 minutes. After washing in PBS, sections were incubated at 65°C for 30 minutes and then transferred to a blocking solution for 1 hour at room temperature. Sections were then stained with an anti-myc antibody (Santa Cruz, SC-40) and placed at 4°C overnight in the dark. Next day, sections were washed and probed with an anti-mouse secondary antibody conjugated to alkaline phosphatase (Jackson, 115-055-062) and incubated for 2 hours at room temperature. To develop the phosphatase signal, labeled sections were incubated for 1 hour in NBT/BCIP solution (Roche, 11681451001) and then washed and mounted in Prolong Gold mountant.

## QUANTIFICATIONS AND STATISTICAL ANALYSIS

### Analysis of *in vivo two-photon* spine dynamics

All spine image analysis was measured manually on the raw image stacks using ImageJ software (NIH ImageJ), blind to the experimental conditions. Only dendritic segments of high-quality image stacks were used for quantification to ensure that tissue rotation or movements between imaging sessions did not affect the identification of dendritic spines. All spines with saturated pixels were excluded. The spines and dendrites of each animal were pooled to calculate a single value of spine density, formation rate, elimination rate, and spine survival for each animal (per day). Spine formation rate: number of newly formed spines on image session S_X_ / total number of spines on session S_X−1_, elimination rate: number of spines missing on session S_X_ (present on session S_X−1_, but not S_X_) / total number of spines on session S_X−1_. Spine survival was quantified as the percentage of spines from session S_X_ that remained present on some future session S_X+ΔX_. Data on spine dynamics throughout this study are presented as mean ± s.e.m.

### Statistics

Repeated measurements (e.g. reaching task success across days, spine dynamics across days, etc.) were analyzed using 2-way repeated measures ANOVA with post-hoc tests. All two-sample comparisons (e.g. spine density comparisons, mEPSC frequency comparisons, etc.) were analyzed with nonparametric tests (Mann-Whitney or Wilcoxon). All correlations were calculated as Pearson correlation coefficient (r) with accompanying p-values. Unless otherwise specified, data is presented as mean ± SEM (standard error of the mean), with all statistical tests, statistical significance values, and sample sizes described in the figure legends.

## DATA AND CODE AVAILABILITY

*In vivo* imaging datasets, electrophysiology datasets, and supporting code for analysis have not been deposited in a public repository but are available from the corresponding authors upon request.

### SUPPLEMENTARY FIGURE LEGENDS

**Figure S1 (related to Figure 1).**
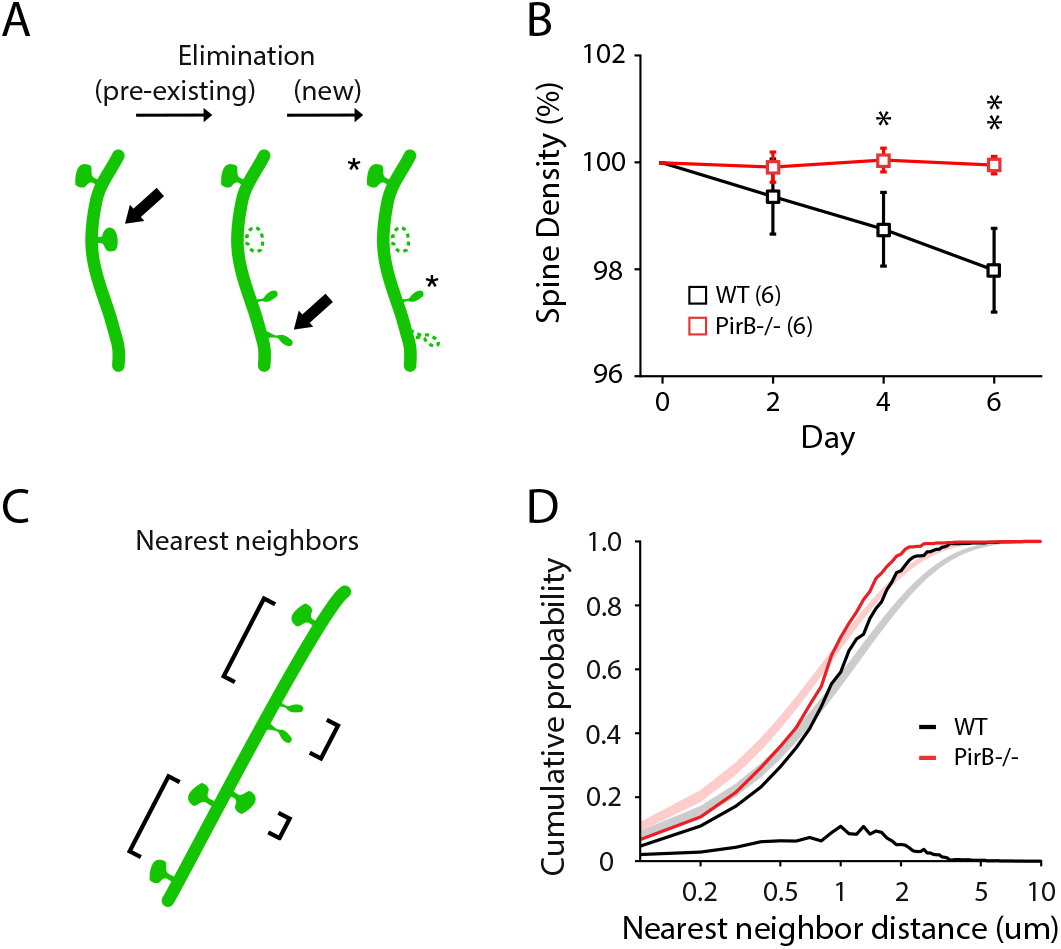
(A) Depiction of spine elimination involving pre-existing spines (those existing upon the first imaging session) vs. new spines (those formed any day starting the second imaging session). (B) Spine density (normalized to the first imaging session) declines significantly less over days in PirB−/− mice (WT: n = 6; PirB−/−: n = 6; day 2, WT: 99.37 ± 0.29%, PirB−/−: 99.92 ± 0.12%, p = 0.8182; day 4, WT: 98.75 ± 0.29%, PirB−/−: 100.05 ± 0.09%, p = 0.0455; day 6, WT: 97.98 ± 0.32%, PirB−/−: 99.96 ± 0.07%, p = 0.0043, Mann-Whitney). (C) For each spine, the distance to its nearest neighbor (NN) was identified in order to quantify spine clustering along dendrites. (D) Cumulative distributions were generated for NNs across all spines of both genotypes (solid lines). NNs drawn from shuffled data were used to generate expected random distributions (shaded regions). Both WT and PirB−/− mice spines had significantly more clustering than expected, with more spines represented in the 1-3μm range (WT: n = 3158 spines / 7 mice; p = 9.3e-19; PirB−/−: n = 3986 spines / 8 mice; p = 2.1e-26, 2-sample Kolmogorov-Smirnov). Furthermore, spine NNs in PirB−/− were significantly smaller than those of WT (p = 1.0e-18, 2-sample Kolmogorov-Smirnov), with a greater frequency of spines with 0.2-2.0μm separation (bottom black line).

**Figure S2 (related to Figure 2).**
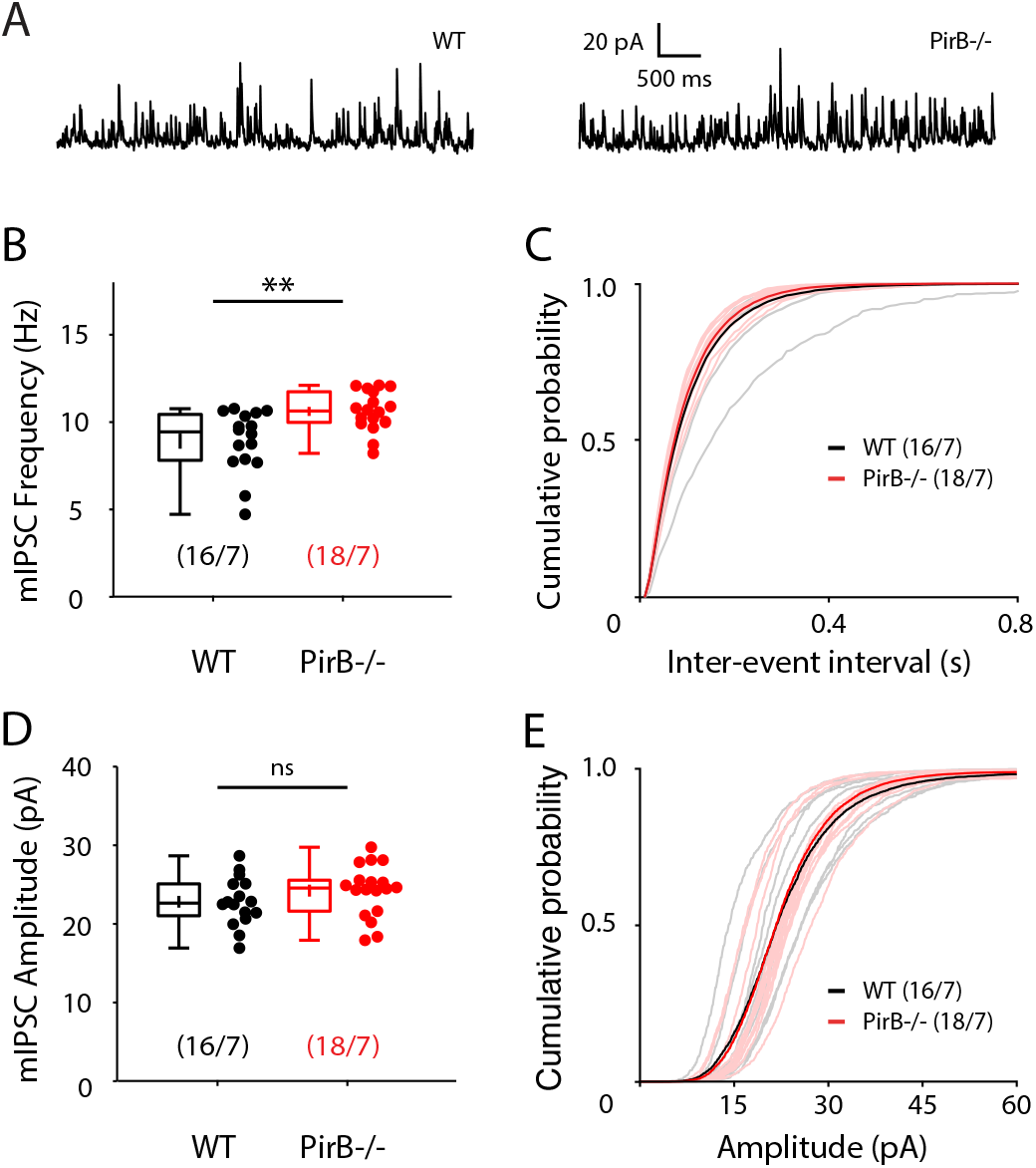
(A) Representative recordings of mIPSCs from M1 layer 5 pyramidal neurons WT and PirB−/− mice. Tetrodotoxin (1μM), NBQX (10μM), and R-CPP (10μM) were included in the ACSF. (B, C) The frequency of mIPSCs was significantly increased (decreased inter-event interval) in PirB−/− mice, compared to WT littermates (WT: 8.910 ± 0.431; n = 16 cells / 7 mice; PirB−/−: 10.611 ± 0.259; n = 18 cells / 7 mice; p = 0.0028, Mann-Whitney). (D, E) mIPSC amplitudes were not significantly different between WT and PirB−/− mice (WT: 22.816 ± 0.748; n = 16 cells / 7 mice; PirB−/−: 24.228 ± 0.757; n = 18 cells / 7 mice; p = 0.2079, Mann-Whitney). (B-E) Circles represent individual mice. Box plots and mean ± s.e.m. are shown. ** p < 0.01; ns: non-significant.

**Figure S3 (related to Figure 3).**
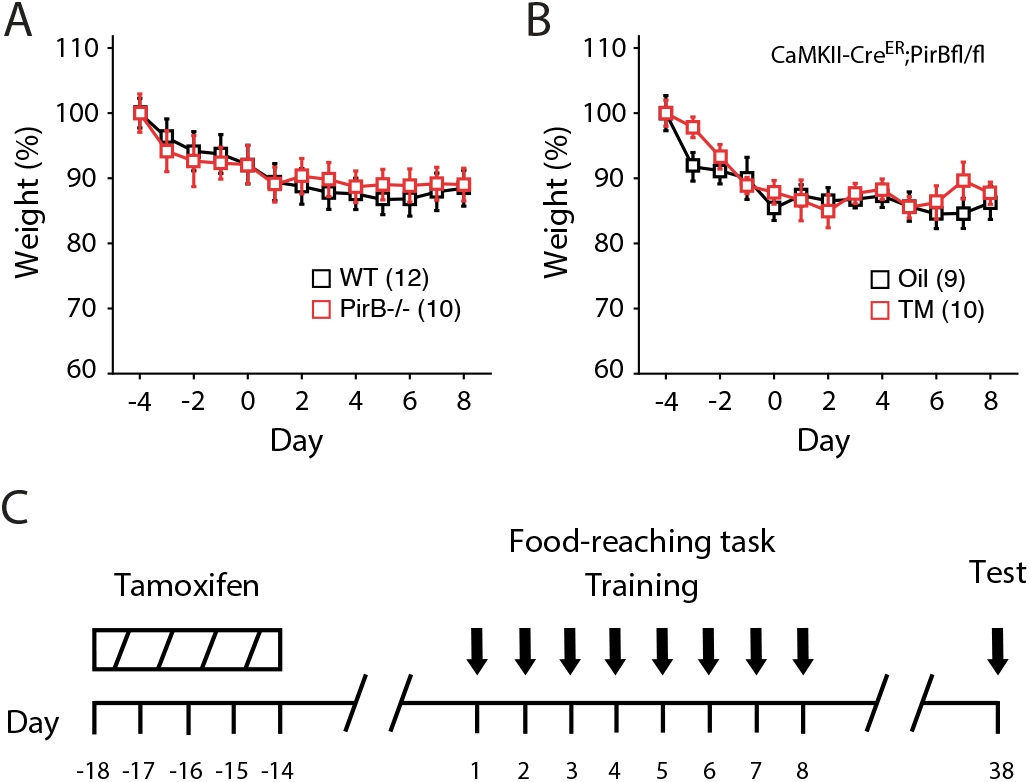
(A) Summary of WT and PirB−/− animal weights throughout the behavioral paradigm. Food restriction began at day −4 and the first training session began day 1. Both WT and PirB−/− mice rapidly dropped to 90% starting body weights, and there were no significant fluctuations in body weight throughout training (WT: n = 12; PirB−/−: n = 10; p = 0.907, 2-way repeated measures ANOVA). (B) Baseline-normalized weights for tamoxifen-injected and control animals. There were no significant fluctuations in body weight throughout training (oil: n = 9; tamoxifen: n = 10; p = 0.700, 2-way repeated measures ANOVA). (C) Experimental timeline for tamoxifen-injected animal training. CaMKII-CreER;PirB^fl/fl^ mice received 5 consecutive days of tamoxifen (4 mg / day, or oil vehicle control) such that the last injection was ~1 week before food restriction and ~2 weeks before the first training day. Animals were retested 30 days following initial training in order to assess long-term memory of the newly acquired motor skill. (A, B) Squares depict mean (± s.e.m.) within each genotype / condition.

**Figure S4 (related to Figure 4).**
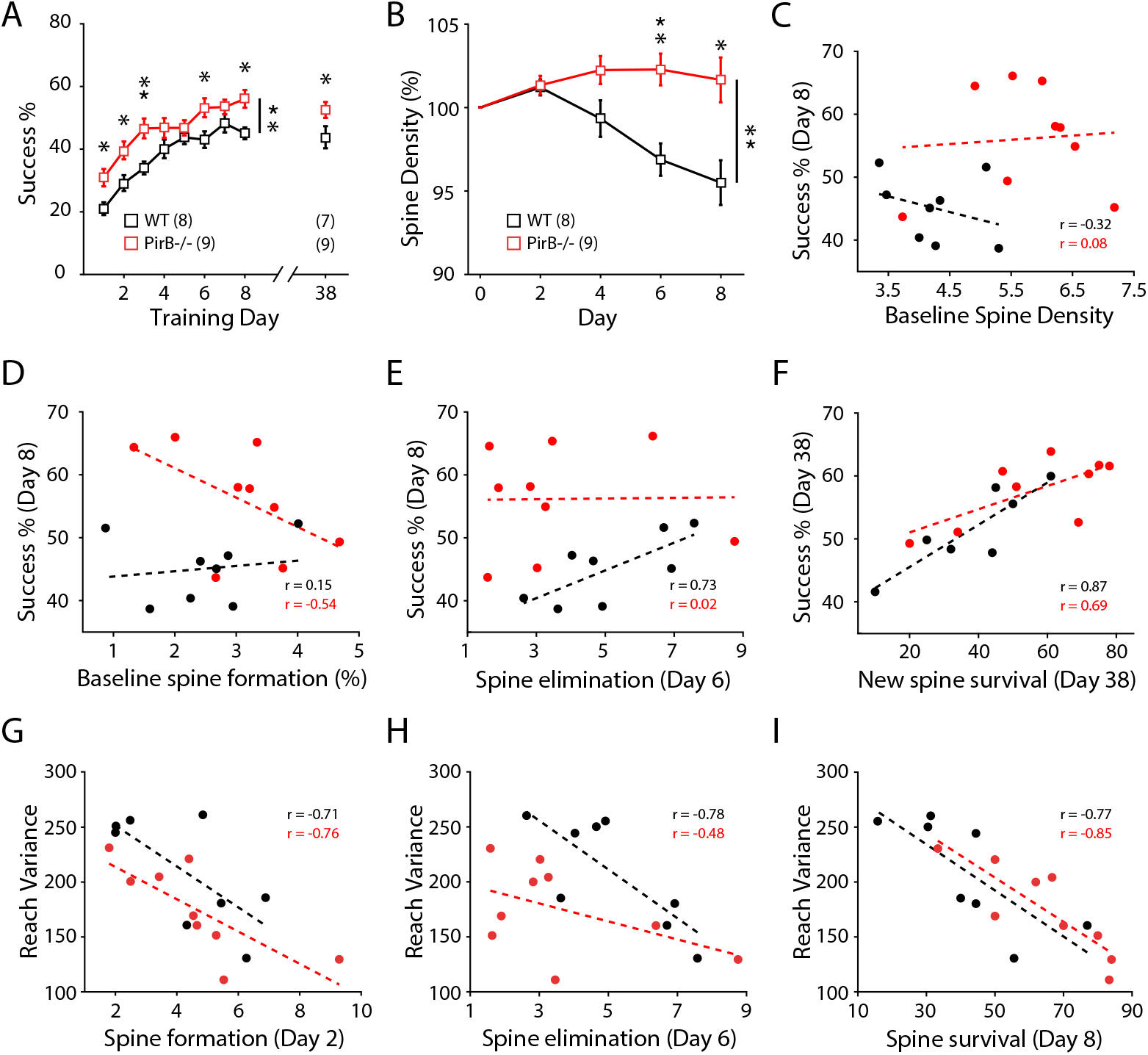
(A) Mice were trained for 8 days followed by retesting at day 38. As before, PirB−/− mice performed the reaching task better compared to WT during 8 training days (WT: n = 8; PirB−/−: n = 9; p = 0.001, 2-way repeated measures ANOVA) and again when re-assessed 30 days after training (WT: X ± X%; n = 7; PirB−/−: X ± X%; n = 9; p = 0.031, Mann-Whitney). (B) Spine density throughout training (normalized to the last baseline day) is significantly elevated in PirB−/− mice (WT: n = 8; PirB−/−: n = 9; p = 0.001, 2-way repeated measures ANOVA). Individual day statistics (day 2, WT: 101.21 ± 0.17%, PirB−/−: 101.34 ± 0.19%, p = 0.9996; day 4, WT: 99.34 ± 0.39%, PirB−/−: 102.25 ± 0.28%, p = 0.1983; day 6, WT: 96.88 ± 0.34%, PirB−/−: 102.28 ± 0.32%, p = 0.0049; day 8, WT: 95.59 ± 0.47%, PirB−/−: 101.66 ± 0.45%, p = 0.0216, Sidak posthoc multiple comparisons). (C) Correlation between baseline spine density and last day performance on reaching task (WT: r = −0.3193; n = 8; p = 0.4407, Pearson correlation; PirB−/−: r = 0.0798; n = 9; p = 0.8382, Pearson correlation). (D) Correlation between baseline spine formation and last day performance on reaching task (WT: r = 0.1490; n = 8; p = 0.7247, Pearson correlation; PirB−/−: r = −0.5435; n = 9; p = 0.1304, Pearson correlation). (E) Correlation between day 6 spine elimination and last day performance on reaching task (WT: r = 0.7255; n = 8; p = 0.0416, Pearson correlation; PirB−/−: r = 0.0154; n = 9; p = 0.9687, Pearson correlation). (F) Correlation between new spine survival on day 38 and day 38 performance on reaching task (WT: r = 0.8746; n = 7; p = 0.0100, Pearson correlation; PirB−/−: r = 0.6919; n = 9; p = 0.0389, Pearson correlation). (G) Correlation between spine formation on day 2 and last day reach trajectory variance (WT: r = −0.7110; n = 8; p = 0.0480, Pearson correlation; PirB−/−: r = −0.7620; n = 9; p = 0.0169, Pearson correlation). (H) Correlation between spine elimination on day 6 and last day reach trajectory variance (WT: r = −0.7778; n = 8; p = 0.0231, Pearson correlation; PirB−/−: r = −0.4760; n = 9; p = 0.1953, Pearson correlation). (I) Correlation between new spine survival on day 8 and last day reach trajectory variance (WT: r = −0.7665; n = 8; p = 0.0265, Pearson correlation; PirB−/−: r = −0.8461; n = 9; p = 0.0040, Pearson correlation). (A-C) Squares depict mean (± s.e.m.) within each genotype. * p < 0.05, ** p < 0.01. (D-I) Circles depict individual animals, dotted lines depict linear fit of each correlation.

**Figure S5 (related to Figure 6).**
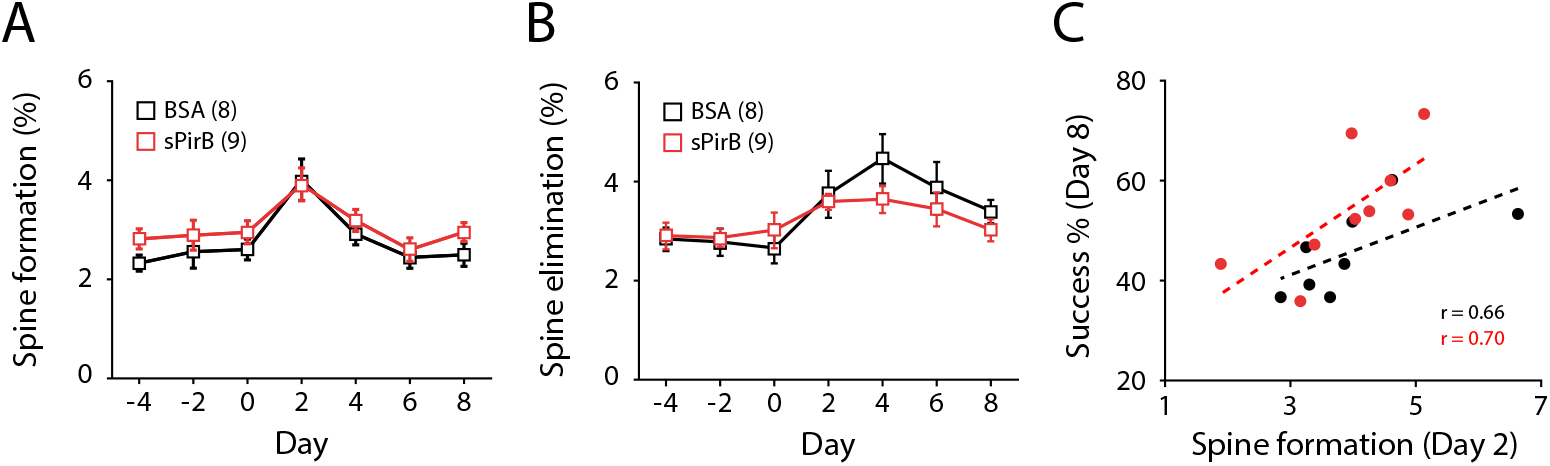
(A) Spine formation rates throughout training in BSA or sPirB infused mice were not significantly different (p = 0.1166, 2-way repeated measures ANOVA). (B) Spine elimination rates throughout training in BSA or sPirB infused mice were not significantly different (p = 0.3813, 2-way repeated measures ANOVA). (C) Correlation between new spine formation on day 2 and last day performance on reaching task (WT: r = 0.6583; n = 8; p = 0.0759, Pearson correlation; PirB−/−: r = 0.6978; n = 9; p = 0.0366, Pearson correlation). (A, B) Squares depict mean (± s.e.m.) within each genotype. (C) Circles depict individual animals, dotted lines depict linear fit of each correlation.

